# Stepwise gating of the Sec61 protein-conducting channel by Sec63 and Sec62

**DOI:** 10.1101/2020.12.14.422728

**Authors:** Samuel Itskanov, Eunyong Park

## Abstract

The universally conserved Sec61/SecY channel mediates transport of many newly synthesized polypeptides across membranes, an essential step in protein secretion and membrane protein integration^1-5^. The channel has two gating mechanisms—a lipid-facing lateral gate, through which hydrophobic signal sequences or transmembrane helices (TMs) are released into the membrane, and a vertical gate, called the plug, which regulates the water-filled pore required for translocation of hydrophilic polypeptide segments^6^. Currently, how these gates are controlled and how they regulate the translocation process remain poorly understood. Here, by analyzing cryo-electron microscopy (cryo-EM) structures of several variants of the eukaryotic post-translational translocation complex Sec61-Sec62-Sec63, we reveal discrete gating steps of Sec61 and the mechanism by which Sec62 and Sec63 induce these gating events. We show that Sec62 forms a V-shaped structure in front of the lateral gate to fully open both gates of Sec61. Without Sec62, the lateral gate opening narrows, and the vertical pore becomes closed by the plug, rendering the channel inactive. We further show that the lateral gate is opened first by interactions between Sec61 and Sec63 in both cytosolic and luminal domains, a simultaneous disruption of which fully closes the channel. Our study defines the function of Sec62 and illuminates how Sec63 and Sec62 work together in a hierarchical manner to activate the Sec61 channel for post-translational translocation.

## Main text

In all organisms, about one third of proteins are transported across or integrated into a membrane upon synthesis by the ribosome. This translocation process mainly occurs in the endoplasmic reticulum (ER) membrane of eukaryotes or the plasma membrane of prokaryotes, mediated by the conserved heterotrimeric protein-conducting channel called the Sec61 (SecY in prokaryotes) complex^1-5^. The main α subunit of the channel, comprised of ten TMs, forms an hourglass-shaped cavity, through which polypeptide chains are transported, whereas the small β and γ subunits peripherally associate with the α subunit in the membrane^6^. Previous structures of Sec61/SecY showed that in the idle state the pore is blocked in the ER luminal (or extracellular) funnel by the plug domain—a structure formed by a segment immediately following TM1 of the α subunit^6-9^, whereas in translocating states the plug moves away^10-12^. Sec61/SecY can also release polypeptides to the lipid phase through a lateral gate formed between TM2 and TM7 of the α subunit, a process required for recognition of hydrophobic targeting signals of soluble secretory proteins and integration of membrane proteins.

The Sec61/SecY channel alone is inactive and thus must associate with partners to enable translocation. In the co-translational mode, common in both prokaryotes and eukaryotes, The channel directly docks with the ribosome-nascent-chain complex^11,13,14^. Many secretory proteins are targeted to the channel post-translationally after their release from the ribosome^15-20^. In eukaryotes, post-translational translocation is enabled by association between the Sec61 complex and the two essential integral membrane proteins Sec62 and Sec63, forming a machinery called the Sec complex^21-24^. In fungal species, the Sec complex also contains the additional nonessential proteins Sec71 and Sec72, which are bound to Sec63 in the cytosol. Sec63 recruits the ER luminal Hsp70 ATPase BiP to the complex to power translocation^25,26^. In bacteria, a single cytosolic ATPase called SecA binds to the SecY complex to drive post-translational translocation^10,12,19,20,27^.

Recently, two cryo-EM studies reported structures of the Sec complex from *Saccharomyces cerevisiae* at ~4 Å resolution^28,29^, which suggested a putative role of Sec63 in activating the Sec61 channel for translocation. However, in both structures, Sec62 was barely visible, and thus its function remains elusive despite its essentiality. Furthermore, the two structures displayed noticeable conformational differences in Sec61, despite essentially identical specimen compositions. Most notably, in one structure the pore is blocked by the plug^29^, whereas in the other structure the plug is displaced leaving the pore open^28^. Although deemed important given the role of the plug in channel gating, the cause of this difference remains a puzzle. Finally, although Sec63 has been suggested to open the lateral gate of Sec61^28,29^, the mechanism of opening remains speculative without structures of mutants and other conformations. Thus, whether and how Sec62 and Sec63 regulate gating of Sec61 are poorly understood. Addressing these issues is essential for our understanding of eukaryotic post-translational translocation and the mechanism of the Sec61/SecY channel in general. Mutations in Sec61 and Sec63 have been implicated in hyperuricemic nephropathy^30^, diabetes^31^, and polycystic liver disease^32^.

### Cryo-EM analysis of two fungal Sec complexes

To determine how the gates are regulated in the Sec complex, we first analyzed a large cryo-EM dataset of the wildtype (WT) Sec complex from *S. cerevisiae* (*Sc*Sec), which yielded three structures at 3.1-3.2-Å resolutions with distinct conformations (Fig. 1a,b, Table 1, and Extended Data Fig. 1). While reconstruction from approximately 1 million particles yielded a 3.0-Å-resolution consensus map (Extended Data Fig. 1b-d), we found that the particle set contained subpopulations lacking Sec62 or Sec71-Sec72, despite apparent sample homogeneity (Extended Data Fig. 1a). We therefore performed additional three-dimensional (3D) classifications to separate particles with and without Sec62 (referred to as Sec62+ and Sec62-) (Extended Data Fig. 1b,e). Furthermore, the Sec62+ class could be further separated into two distinct subclasses (referred to as C1 and C2), which show notable conformational differences in Sec62, the lateral gate, and the plug (Fig 1b and Extended Data Fig. 1f,g; see below). Although an atomic model for Sec62 could not be built due to insufficient local resolution, the classification significantly improved Sec62 features, enabling unambiguous assignment of individual domains (Fig. 1d).

**Figure 1.**
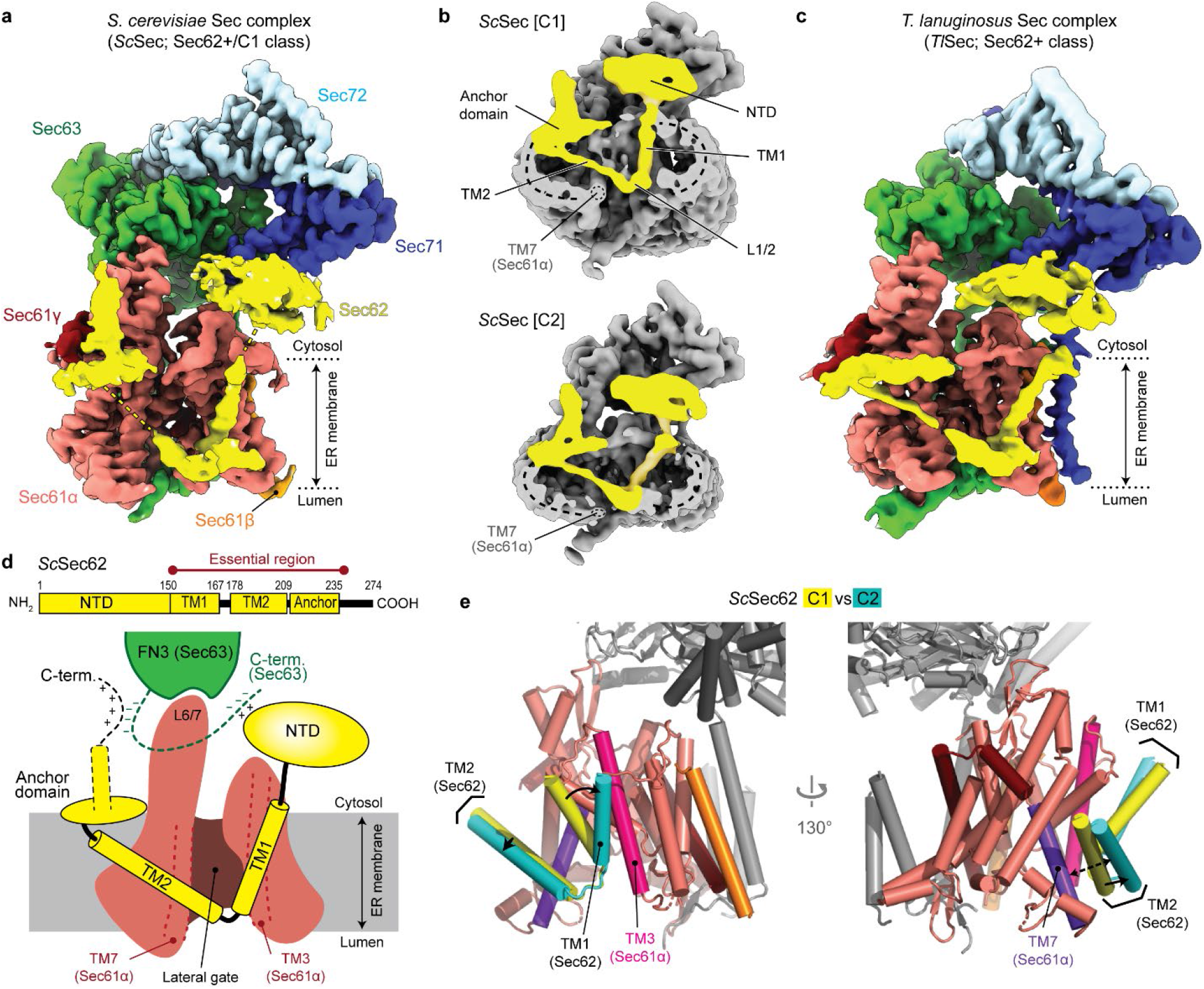
Cryo-EM analysis of fungal Sec complexes and the structure of Sec62. **a**, The 3.1-Å-resolution cryo-EM reconstruction of the yeast Sec complex (front view into the lateral gate). Yellow dash lines indicate the connections that are visible at a lower contour level (see panel b). In yeast nomenclature, the α, β, and γ subunits of the Sec61 complex are called as Sec61p, Sbh1p, and Sss1p, respectively. **b**, Cutaway views showing Sec62 (yellow). Shown are 6-Å-lowpass-filtered C1 (upper panel; a tilted view from the ER lumen) and C2 (lower panel; front view) maps. Dashed line, detergent micelle. **c**, The 3.8-Å-resolution reconstruction of the *T. lanuginosus* Sec complex (the consensus Sec62+ map). **d**, Domain organization of Sec62. Previous studies suggest an interaction between the NTD of Sec62 and the C-terminal tail of Sec63 (ref. ^33,42^). In addition, based on the proximity, the C-terminal tails of Sec62 and Sec63 may also interact with each other through an electrostatic interaction. **e**, Interactions between the Sec62 TMs and lateral gate. Dashed arrow, a gap between Sec61α TM7 and Sec62 TM2 in the C2 conformation. The color scheme for Sec61 is the same as in a. Sec63, Sec71, and Sec72 are in grey.

**Table 1.** Cryo-EM data collection, refinement and validation statistics of wildtype *Sc*Sec and wildtype and mutant *TlSec* complexes

| | Wildtype *ScSec* | | | Wildtype *TtSec* | | | ΔSec62<br>*TtSec* | Δanchor<br>*TtSec* |
| --- | --- | --- | --- | --- | --- | --- | --- | --- |
|  | Sec62– | Sec62+/C1 | Sec62+/C2 | Sec62– | Plug-<br>open | Plug-<br>closed |  |  |
| EM Databank<br>accession code | EMD-22770 | EMD-22771 | EMD-22772 | EMD-22773 | EMD-22774 | EMD-22775 | EMD-22776 | EMD-22777 |
| PDB accession code | 7KAH | 7KAI | 7KAJ | 7KAK | 7KAL | 7KAM | 7KAN | N/A |
| **Data collection** |  |  |  |  |  |  |  |  |
| Magnification | 64,000x |  |  | 36,000x |  |  | 36,000x | 64,000x |
| Voltage (kV) | 300 |  |  | 200 |  |  | 200 | 300 |
| Electron exposure (e<sup>−</sup><br>/Å<sup>2</sup>) | 49.1 |  |  | 50.0 |  |  | 50.0 | 49.1 |
| Defocus range (μm) | -0.8 to -2.5 |  |  | -0.6 to -2.4 |  |  | -0.9 to -2.2 | -0.7 to -2.9 |
| Pixel size (Å) | 1.19 |  |  | 1.14 |  |  | 1.14 | 1.19 |
| **Processing** |  |  |  |  |  |  |  |  |
| Symmetry imposed | C1 | C1 | C1 | C1 | C1 | C1 | C1 | C1 |
| Initial particle images<br>(no.) | 2,686,839 |  |  | 1,632,659 |  |  | 546,712 | 229,825 |
| Final particle images<br>(no.) | 391,885 | 193,263 | 193,661 | 155,601 | 114,704 | 143,227 | 222,047 | 76,726 |
| Map resolution (Å) | 3.1 | 3.2 | 3.1 | 3.9 | 4.0 | 3.8 | 3.7 | 4.4 |
| FSC threshold | 0.143 | 0.143 | 0.143 | 0.143 | 0.143 | 0.143 | 0.143 | 0.143 |
| Map resolution range<br>(Å) | 2.6 –<br>11 | 2.8 –<br>12 | 2.7 –<br>12 | 3.4 –<br>13 | 3.3 –<br>13 | 3.3 –<br>12 | 3.3 –<br>12 | 3.7 –<br>14 |
| **Refinement** |  |  |  |  |  |  |  |  |
| Initial model used<br>(PDB code) | 6N3Q | 7KAH | 7KAH | 7KAN | 7KAN | 7KAN | 6N3Q | - |
| Model resolution (Å) | 3.2 | 3.3 | 3.3 | 4.1 | 4.2 | 4.0 | 4.0 | - |
| FSC threshold | 0.5 | 0.5 | 0.5 | 0.5 | 0.5 | 0.5 | 0.5 | - |
| Map sharpening *B*<br>factor (Å<sup>2</sup>) | 86.6 | 80.8 | 75.9 | 110.3 | 90.7 | 105.2 | 127.8 | - |
| **Model composition** |  |  |  |  |  |  |  |  |
| Nonhydrogen atoms | 10,495 | 10,718 | 10,712 | 10,438 | 10,794 | 10,921 | 10,661 | - |
| Protein residues | 1,349 | 1,399 | 1,399 | 1,371 | 1,429 | 1,445 | 1,371 | - |
| Ligands | - | - | - | - | - | - | 2 (PC2) | - |
| *B* factors, average (Å<sup>2</sup>) | 73 | 61 | 58 | 117 | 126 | 74 | 30 | - |
| **R.m.s. deviations** |  |  |  |  |  |  |  |  |
| Bond lengths (Å) | 0.003 | 0.003 | 0.003 | 0.002 | 0.002 | 0.003 | 0.003 | - |
| Bond angles (°) | 0.522 | 0.508 | 0.513 | 0.524 | 0.489 | 0.521 | 0.623 | - |
| **Validation** |  |  |  |  |  |  |  |  |
| MolProbity score | 1.43 | 1.42 | 1.33 | 1.51 | 1.42 | 1.48 | 1.55 | - |
| Clashscore | 4.61 | 4.14 | 3.87 | 6.33 | 5.62 | 5.60 | 6.18 | - |
| Poor rotamers (%) | 0 | 0 | 0 | 0 | 0 | 0 | 0 | - |
| **Ramachandran plot** |  |  |  |  |  |  |  |  |
| Favored (%) | 96.83 | 96.58 | 97.01 | 97.09 | 97.42 | 96.96 | 96.72 | - |
| Allowed (%) | 3.17 | 3.42 | 2.99 | 2.91 | 2.58 | 3.04 | 3.28 | - |
| Disallowed (%) | 0 | 0 | 0 | 0 | 0 | 0 | 0 | - |

To gain insights into structural and mechanistic conservation across species, we also determined structures of the Sec complex from the thermophilic fungus *Thermomyces lanuginosus* (*Tl*Sec) at overall resolution of 3.6 to 3.9 Å (Fig. 1c, Table 1, and Extended Data Fig. 2). *Tl*Sec classes with and without Sec62 closely resemble the *Sc*Sec [C2] and [Sec62-] structures, respectively (brackets denote classes). We could not isolate a C1-equivalent class from the *Tl*Sec dataset perhaps because the specimen freezing condition (4°C) might have biased the conformation distribution of this thermophilic complex towards C2. The structure of *Tl*Sec[Sec62-] is essentially identical to a separately determined structure of a mutant complex completely lacking Sec62 (ΔSec62 *Tl*Sec) (Extended Data Fig. 2g-i). Importantly, the domain arrangement of *Tl*Sec62 is the same as that of *Sc*Sec62 despite ~30% overall sequence identity (Fig. 1c and Extended Data Fig. 3). Compared to *Sc*Sec62, *Tl*Sec62 is better resolved such that we could register amino acids to its TM1.

### Sec62 forms a V-shaped structure

Sec62 consists of a cytosolic, globular N-terminal domain (NTD), two TMs (TM1 and TM2) connected by a short ER luminal loop (L1/2), and a cytosolic C-terminal segment (Fig. 1d). Functionally essential regions have previously been mapped to the two TMs and a segment of ~30 amino acids immediately following TM2 (ref. ^33^). The TMs of Sec62 are arranged as a V shape in front of the lateral gate with L1/2 directed to the lateral gate opening (Fig. 1a-d). The contact with the channel is mainly formed by an interaction of Sec62-TM1 with the TM3 and N-terminal segment of Sec61α.

Following TM2, Sec62 contains an oval-shaped structure lying flat on the membrane interface (Fig. 1a-d, and Extended Data Fig 4a-b). This amphipathic structure, which we termed the anchor domain, is most likely formed by an ~20-residue-long conserved segment within the abovementioned 30 amino acids, and is rich in hydrophobic amino acids (Extended Data Fig. 3). While single-point mutations of these hydrophobic residues caused no growth defect, alanine substitutions of three consecutive residues in positions 215-220 were lethal (Extended Data Fig. 4d), suggesting that decreased hydrophobicity interrupts its functionally essential interaction with the membrane. The structure of a *Tl*Sec mutant with the anchor domain replaced with a glycine/serine linker (Δanchor *Tl*Sec) showed virtually no visible Sec62 features (Extended Data Fig. 4e,f), suggesting Sec62 becomes too mobile without the domain. Taken together, these observations suggest that the function of the anchor domain is to properly position the V-shaped TMs of Sec62 at the lateral gate.

The revealed position and topology of Sec62 raise an important question about how the channel would engage with substrate polypeptides. During the initial stage of post-translational translocation, a substrate polypeptide is expected to insert into the channel as a loop with both its N- and C-termini exposed to the cytosol^34^ (Extended Data Fig. 5a). While the N-terminal signal sequence may sit initially at the lateral gate as seen in structures of mammalian co-translational and bacterial post-translational complexes^10-12^, later it must engage with the signal peptidase for cleavage^35^. However, the presence of Sec62 would pose a problem because L1/2 of Sec62 would block the release of the signal sequence from the lateral gate. The answer may be provided by a conformational transition from C1 to C2 as visualized in the *Sc*Sec structure (Fig. 1e). While in both structures the seam between the Sec62-TM1 and Sec61α-TM3 is tight, a sufficient gap is formed on the other side of the lateral gate between the Sec62-TM2 and Sec61α-TM7 in the *Sc*Sec[C2] structure. A similar gap also exists in the *Tl*Sec structures (Extended Data Fig. 5b). Thus, the signal sequence of the substrate likely exits through the gap transiently formed between Sec62-TM2 and Sec61α-TM7 during translocation.

### Sec62 regulates the gates of Sec61

Three distinct classes of *Sc*Sec (i.e., C1, C2, and Sec62-) showed notable conformational differences in the lateral gate (Fig. 2a, Supplementary Movie 1). Although open in all three structures, the extent of the lateral gate opening varies on the ER luminal side, with C1 most open and Sec62-least open. The C2 structure, in which Sec62-TM2 is disengaged, is open to an intermediate degree. The movement is mainly mediated by a rigid-body rotation of the TM7, TM8, and the intervening loop (L7/8) of Sec61α (Fig. 2a), which seems to be induced by the interaction between L1/2 of Sec62 and the lateral gate (Fig. 1a-d). Thus, this movement is distinct from the hinge-like motion between the two halves (TM1-5 and TM6-10) of Sec61α which mediates opening of the channel from the fully closed state^6,36-38^.

**Figure 2.**
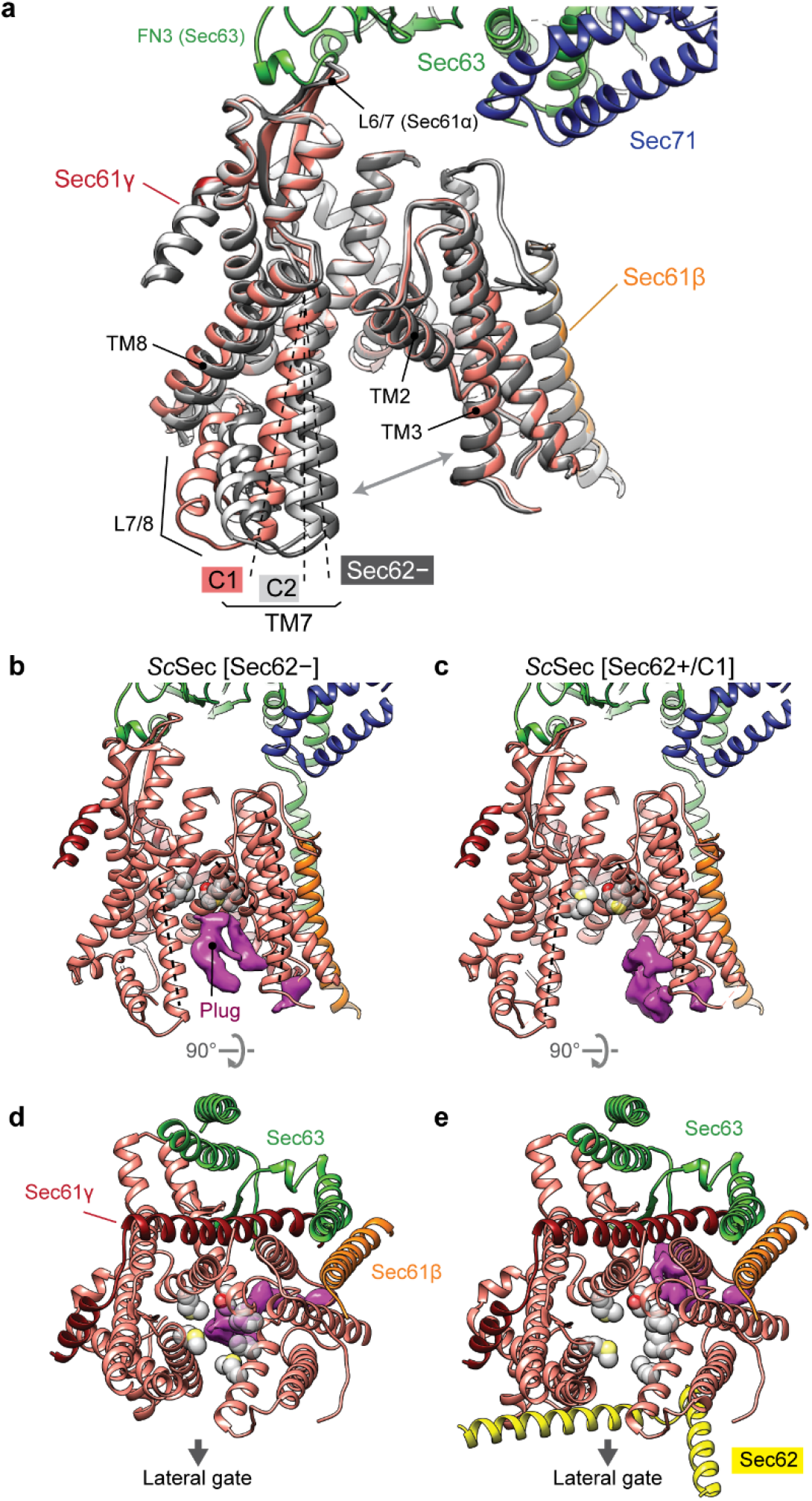
Regulation of the lateral and vertical gates by Sec62. **a**, A comparison of the Sec61 channel conformation between the three *Sc*Sec classes, C1 (in color), C2 (light grey) and Sec62-(dark grey). Dashed lines, TM7 of Sec61α. Grey arrows, the lateral gate. Sec62 is not shown. **b-e**, A comparison of the plug domain (purple density) between Sec62-lacking and -containing *Sc*Sec classes. Grey spheres, pore ring residues. Dashed lines, lateral gate helices (left to right: TM7, TM2, and TM3 of Sec61α). Shown are front views (b and c) and cytosolic views (d and e).

Importantly, the motion of TM7-8 of Sec61α appears to control the position of the plug (Fig. 2 b-e). In *Sc*Sec[Sec62-], the plug is clearly visible immediately below the pore constriction (‘plug-closed’ conformation; Fig. 2 b,d, Extended Data Fig. 6). By contrast, in *Sc*Sec[C1], the plug is displaced to a position near the C-terminus of Sec61γ (‘plug-open’ conformation; Fig. 2 c,e), thus opening the pore. In *Sc*Sec[C2], the plug seems disordered, probably because it takes intermediate positions between the two conformations. Similar observations were also made with the *Tl*Sec structures: compared to the Sec62- and ΔSec62 structures, the Sec62+ structure shows a shifted position of Sec61α TM7-8 as in *Sc*Sec[C2] (Extended Data Fig. 7) and concomitant plug mobilization, where 53% and 42% particles classified into the plug-closed and open conformations, respectively (Extended Data Figs. 2 and 7b,c). The plug displacement is likely caused by the Sec62-induced movement of Sec61α TM7 since the plug interacts with TM7 and L7/8 in the plug-closed conformation^36^ (Extended Data Fig. 7e).

### Partially open Sec61 is inactive

Despite the observed channel gating by Sec62, physiological importance of this role remained unclear. Without Sec62, the lateral gate can still be opened by Sec63. Even though the pore is blocked by the plug, it has been proposed that insertion of a substrate polypeptide would push the plug away^29^. To investigate importance of the Sec62-dependent gating, we sought for mutations affecting Sec61 gating as ΔSec62 does, but independently of Sec62. If the gating function of Sec62 is essential, such mutations would be expected to compromise cell viability.

We first chose to mutate the fibronectin III (FN3) domain of Sec63, which interacts with the cytosolic loop 6/7 (L6/7) of Sec61α (Fig. 3a). L6/7 also provides a major interaction site for the ribosome in cotranslational translocation and the SecA ATPase in bacterial post-translational translocation and thus has been universally implicated in priming or activating the channel^7,8,27,39,40^. We found that none of the FN3 mutants had a growth defect at 30°C (Fig. 3b, left). Only a mild defect was seen at 37°C even with the most severe mutant (FN3mut) (Extended Data Fig. 8b). To understand this unexpectedly weak phenotype, we determined the structure of FN3mut *Sc*Sec (Fig. 3d,e, Table 2, and Extended Data Fig. 8c,d). The structure showed that the FN3 domain was indeed disengaged from L6/7 by the mutation, causing ~10° rotation of Sec61 along the membrane normal (Extended Data Fig. 8e,f). Nonetheless, the lateral gate was still open (Fig. 3d). Importantly, the FN3mut complex still exhibited Sec62-induced TM7 movement and plug mobilization (Fig. 3 d,e), which may explain the near-WT growth phenotype of the mutant.

**Figure 3.**
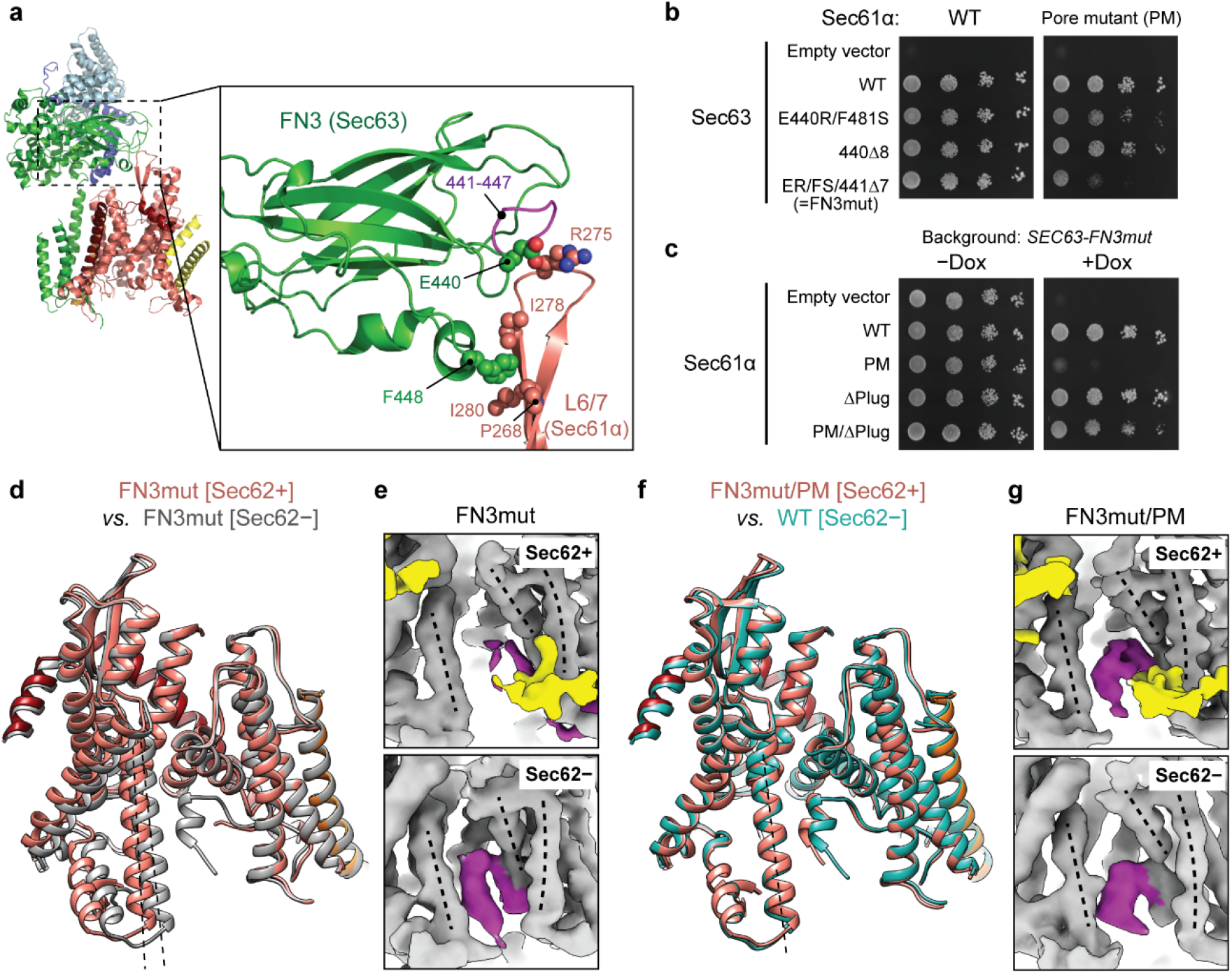
Structural and functional analysis of a gating-defective mutant complex. **a**, The interaction between the FN3 domain of Sec63 and the L6/7 loop of Sec61α. Amino acids involved in the interactions are indicated. **b**, Yeast growth complementation experiments (at 30°C) testing functionality of indicated FN3 mutants of Sec63 in the background of wild-type (left) or pore-mutant Sec61α (right). FN3mut refers to a combination of E440R (ER) and F481S (FS) mutations and a deletion of seven amino acids 441-447 (441Δ7). To repress chromosomal WT Sec63 expression (under a tetracycline promoter), doxycycline was added. Also see Extended Data Fig. 8a. **c**, As in b, but testing for indicated Sec61α mutants in the background of Sec63-FN3mut as a sole Sec63 copy. The addition of doxycycline (Dox) represses chromosomal WT Sec61α expression. **d**, As in Fig. 2a, but with the FN3mut *Sc*Sec structures with and without Sec62. **e**, A comparison of the plug domain (purple density) between the FN3mut *Sc*Sec structures with and without Sec62 (yellow). Dashed lines, lateral gate helices (left to right: TM7, TM2, and TM3 of Sec61α). **f**, As in d, but comparing the Sec62-containing FN3mut/PM structure and the Sec62-class of WT *Sc*Sec. **g**, As in e, but with the FN3mut/PM *Sc*Sec structures.

**Table 2.**
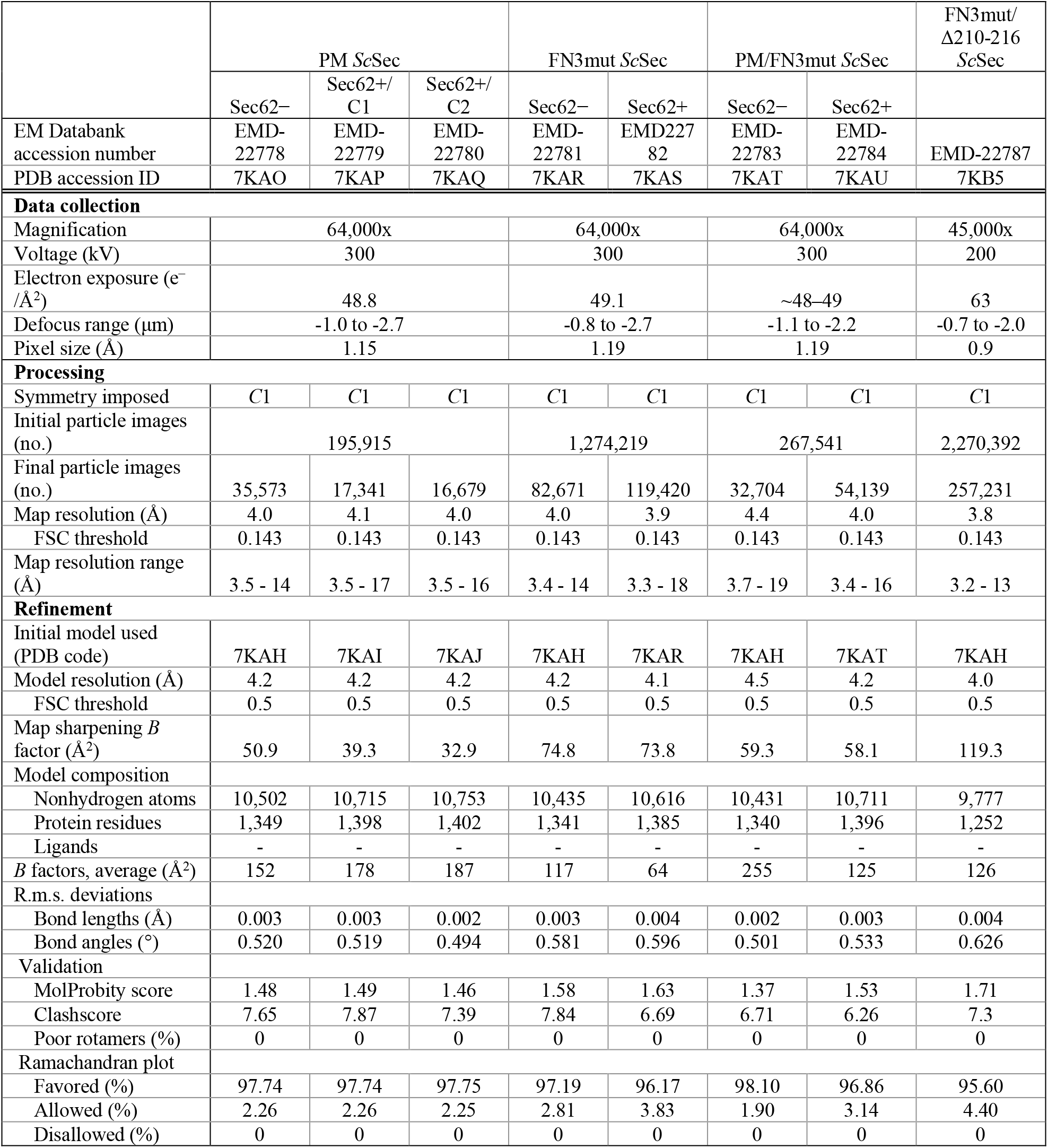
Cryo-EM data collection, refinement and validation statistics of mutant *Sc*Sec complexes

Next, we mutated the pore of Sec61α. In closed SecY structures^6-9^, the aliphatic amino acids lining the pore constriction (called the pore ring residues) make a hydrophobic interaction with the plug. Compared to other species, the pore ring of *Sc*Sec61α appears significantly less hydrophobic^28^. Thus, we reasoned that a mutant with a more hydrophobic pore ring (M90L/T185I/M294I/M450L; collectively denoted PM) might bias the plug towards the closed conformation. In growth complementation assays, PM itself did not affect cell growth, but strikingly, strong synthetic growth impairment was observed when combined with FN3mut (Fig. 3b, right). Importantly, a plug deletion^41^ (ΔPlug) could rescue growth, suggesting that the growth inhibition originates from a gating defect (Fig. 3c). Consistent with this idea, the structures of the combined mutant (FN3mut/PM) showed a strong density of the plug in the closed conformation and no Sec62-dependent movement of lateral gate helices (Fig. 3f,g, and Extended Data Fig. 8g), thereby closely resembling the gating state of *Sc*Sec[Sec62-] despite the presence of Sec62 in front of the lateral gate. On the other hand, PM alone still showed Sec62-mediated movements in the lateral gate and plug, similar to WT (Extended Data Fig. 8h,i). Taken together, these results show that the channel conformation seen in the absence of Sec62 is inactive for post-translational translocation.

### Sec62 clears the lateral gate of lipids

In addition to the role in channel gating, the ΔSec62 *TlSec* structure suggests another function of Sec62— preventing lipids from moving into the lateral gate (Extended Data Fig. 9a-c). In ΔSec62 *Tl*Sec, strong, well-ordered densities of lipid or detergent tails are visible at the lateral gate. The densities are vertically aligned along the hydrophobic groove of the open lateral gate. By contrast, in the Sec62+ structures, only weak fragmented densities were observed. In the cytosolic leaflet, a lipid/detergent molecule seems to be accommodated with an outward rotation of the TM2-3 of Sec61α (Extended Data Fig. 9d). Sec62 may prevent the binding of a lipid by restricting this movement. In the ER luminal leaflet, the L1/2 of Sec62 seems to sterically block lipid binding (Extended Data Fig. 9c). We did not observe a strong lipid/detergent density in the lateral gate of *Sc*Sec[Sec62-], perhaps because of a lower affinity to lipid/detergent. However, one of the previous *Sc*Sec structures^29^, whose conformation resembles the ΔSec62 *Tl*Sec structure, has shown a lipid-like density at the lateral gate and the movement of Sec61α TM2-3 similarly to ΔSec62 *Tl*Sec. Collectively these observations suggest that in the absence of Sec62, lipid molecules may occupy the lateral gate that is opened by Sec63.

### Mechanism of Sec61 gating by Sec63

Our unexpected finding that the FN3-L6/7 interaction is dispensable indicated that there must be another mechanism for Sec63 to open the lateral gate. Besides the FN3 domain, Sec63 forms major contacts with Sec61 through two other parts: TM3, which anchors Sec63 to the Sec61 complex, and a short ER luminal segment (residues 210-216) preceding the TM3, which together with the N-terminal segment of Sec63, interacts with a crevice on the back of the channel (opposite from the lateral gate) (Fig. 4a). We reasoned that the latter interaction might control lateral gating through a lever-like mechanism. In the WT background, replacement of this segment with a glycine/serine linker (Δ210-216) alone did not cause growth inhibition (Fig. 4b). However, when combined with FN3mut, cells did not grow (Fig. 4b).

**Figure 4.**
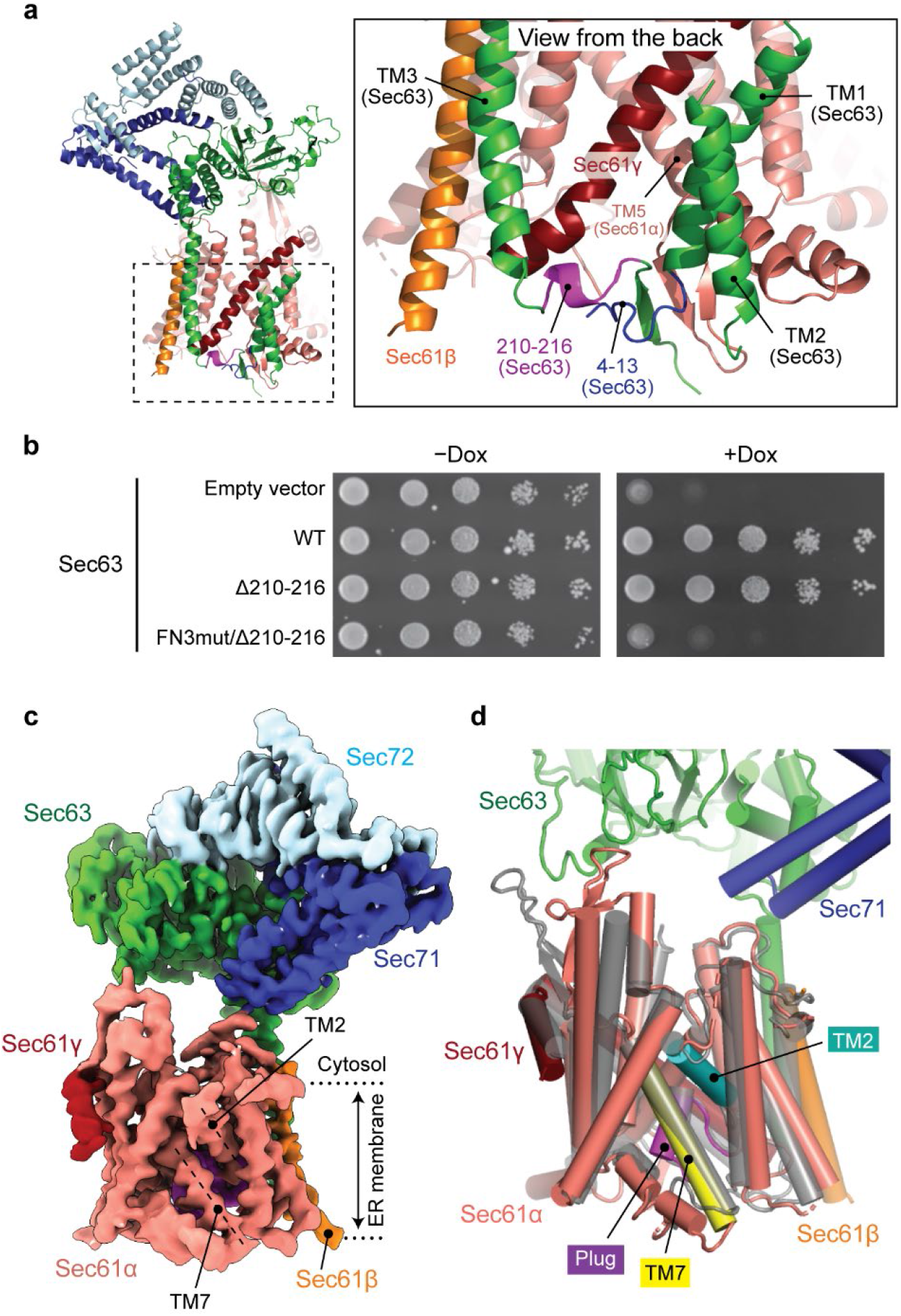
The structure of a fully closed Sec complex. **a**, The interaction between Sec61 and Sec63 in the ER lumen (view from the back). The N-terminal segment (positions 4-13) and the segment preceding TM3 (positions 210-216) of Sec63 are in blue and purple, respectively. **b**, Yeast growth complementation (at 30°C) testing functionality of the indicated Sec63 mutants. The addition of doxycycline (Dox) represses chromosomal WT Sec63 expression. **c**, The 3.8-Å-resolution cryo-EM structure of the *Sc*Sec complex containing FN3mut/Δ210-216 double-mutant Sec63. The lateral gate helices TM2 and TM7 are indicated. **d**, As in c, but showing the atomic model of the Sec61 complex. For comparison, the closed *M. jannaschii* SecY structure (PDB 1RH5; semitransparent grey) is superimposed.

To understand the structural basis of this functional defect, we determined the structure of the FN3mut/Δ210-216 *Sc*Sec complex (Fig. 4c and Extended Data Fig. 10) The structure showed that indeed both the lateral and vertical gates of the Sec61 channel are completely closed, resembling the idle archaeal SecY channel structure^6^ (Fig. 4c and Extended Data Fig. 10). This demonstrates that Sec63 uses both its cytosolic and luminal domains to open the lateral gate in a two-pronged mechanism. The C-terminal cytosolic domain of Sec63 (following TM3) and Sec71-Sec72 are still attached to Sec61 through TM3. However, most of the parts preceding TM3 were invisible due to increased flexibility. Importantly, Sec62 was no longer visible either despite copurification with the complex (Fig. 4c, and Extended Data Fig. 10a). Sec62 is likely associated with Sec63 through an electrostatic interaction with the C-terminal tail of Sec63 (ref.^33,42^) (Fig. 1d), but it seems to no longer bind to the lateral gate due to structural incompatibility with the closed gate. Therefore, the lateral gate must be first opened by Sec63 before Sec62 can activate the channel for protein translocation.

In summary, our study defines the function of Sec62, which has been elusive for three decades since its discovery as an essential component in post-translational translocation^21,22,43^. Once the lateral gate of the Sec61 channel is opened by Sec63, Sec62 fully activates the channel by further mobilizing the plug (Fig. 5). At the same time, Sec62 may keep the open lateral gate clear of lipid molecules, which would likely inhibit signal sequence insertion. The partially open state induced by Sec63 alone seems insufficient for translocation, probably because of a too high energy barrier for a substrate polypeptide to insert into the pore with the plug in place. This contrasts with the co-translational translocation, where the channel opening is thought to be largely mediated by an interaction between the signal sequence (or TM helix) and the lateral gate rather than binding of the ribosome to the channel^2,8,11,14,44^. Many post-translational substrates are known to contain a signal sequence with relatively lower hydrophobicity^15^ and expected to interact more transiently with Sec61 during initial insertion since they are not tethered to the ribosome as in the co-translational mode. These features of post-translational substrates may require both lateral and vertical gates of the channel to be pre-opened for insertion. Our structural analysis shows how Sec63 and Sec62 open the gates of the Sec61 channel in a stepwise fashion to activate the channel. Given the high degree of sequence conservation of these components, the gating mechanism we discovered here is likely conserved across all eukaryotic species.

**Figure 5.**
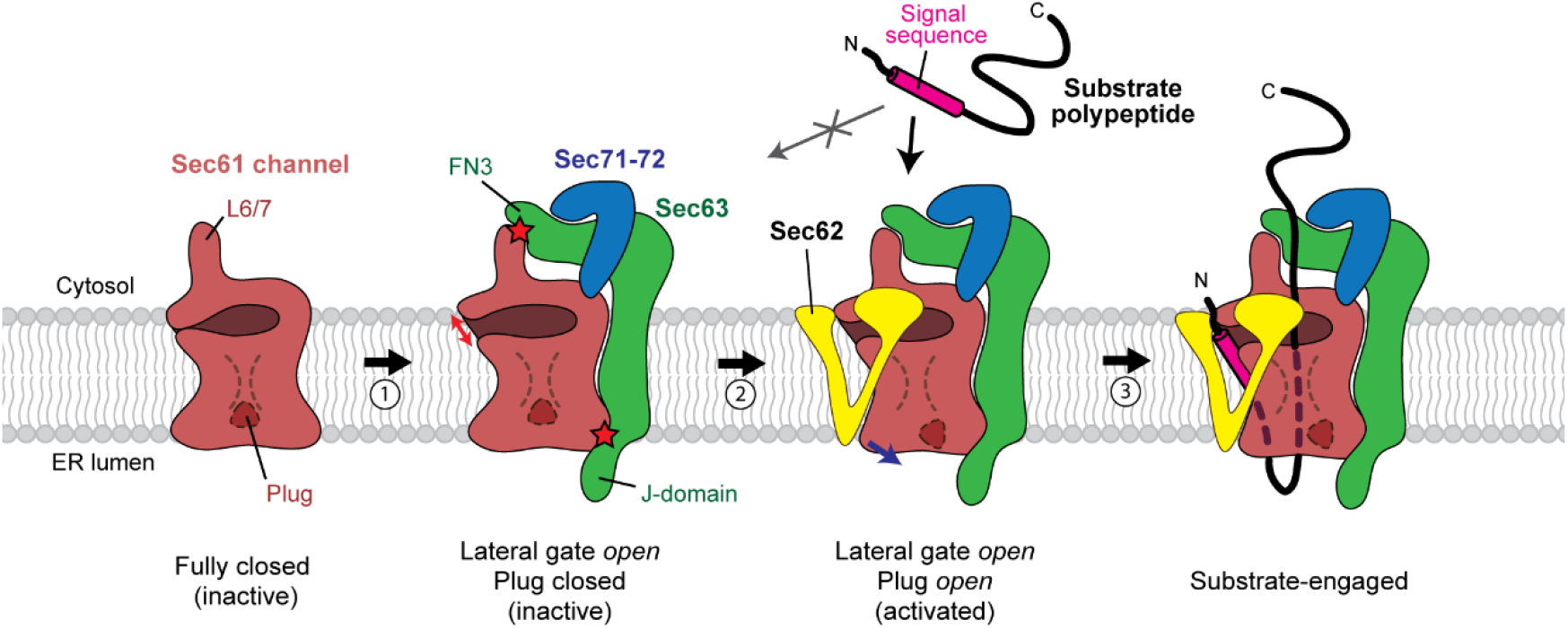
A model for activation of the Sec61 channel by Sec62 and Sec63. The Sec61 channel alone assumes a fully closed conformation (the leftmost cartoon). Step 1, association of Sec63 opens the lateral gate (indicated by a red arrow) through interactions with Sec61 in both the cytosol and ER lumen (indicated by red stars). However, the channel in this conformation is inactive due to the plug in a closed state. Step 2, Sec62 interacts with the lateral gate of Sec61 and further opens the lateral gate (blue arrow), which results in opening of the plug. Step 3, a substrate polypeptide inserts into the open pore of the channel as a loop with the signal sequence sitting at the lateral gate.

## Acknowledgments

We thank D. Toso and J. Remis for support for electron microscope operation, and J. Hurley and E. Nogales for critical reading of manuscript. This work was supported by the Vallee Scholars Program (E.P.).

## Author contributions

S.I. prepared samples and performed functional analysis. S.I and E.P collected and analyzed cryo-EM data, built atomic models, interpreted results, and wrote the manuscript; E.P. supervised the project.

## Data and materials availability

The atomic coordinates of the *Saccharomyces cerevisiae* Sec complexes *Sc*Sec[Sec62-], *Sc*Sec[C1], *Sc*Sec[C2], PM *Sc*Sec[Sec62-], PM *Sc*Sec[C1], PM *Sc*Sec[C2], FN3mut *Sc*Sec[Sec62-], FN3mut *Sc*Sec[Sec62+], PM/FN3mut *Sc*Sec[Sec62-], PM/FN3mut *Sc*Sec[Sec62+] and FN3mut/Δ210-216 *Sc*Sec were deposited to the Protein Data Bank under the accession codes: 7KAH, 7KAI, 7KAJ, 7KAO, 7KAP, 7KAQ, 7KAR, 7KAS, 7KAT, 7KAU and 7KB5, respectively. The atomic coordinates of the *Thermomyces lanuginosus* Sec complexes *Tl*Sec[Sec62-], *Tl*Sec[Sec62+/plug-open], *Tl*Sec[Sec62+/plug-closed], and ΔSec62 *Tl*Sec were deposited to the Protein Data Bank under the accession codes: 7KAK, 7KAL, 7KAM, and 7KAN, respectively.

The cryo-EM density maps for *Sc*Sec[consensus], *Sc*Sec[Sec62-], *Sc*Sec[C1], *Sc*Sec[C2], PM *Sc*Sec[Sec62-], PM *Sc*Sec[C1], PM *Sc*Sec[C2], FN3mut *Sc*Sec[Sec62-], FN3mut *Sc*Sec[Sec62+], PM/FN3mut *Sc*Sec[Sec62-], PM/FN3mut *Sc*Sec[Sec62+], FN3mut/Δ210-216 *Sc*Sec, *Tl*Sec[Sec62-], *Tl*Sec[Sec62+/consensus], *Tl*Sec[Sec62+/plug-open], *Tl*Sec[Sec62+/plug-closed], ΔSec62 *Tl*Sec and Δanchor *Tl*Sec were deposited to the Electron Microscopy Data Bank under the accession codes EMD-22785, EMD22770, EMD-22771, EMD-22772, EMD-22778, EMD-22779, EMD-22780, EMD-227781, EMD-227782, EMD-227783, EMD-22784, EMD-22787, EMD-22773, EMD-22786, EMD-22774, EMD-22775, EMD-22776 and EMD-22777, respectively. All yeast strains and plasmids that were generated in this study are available upon request.

---

**Extended Data Figure 1.**
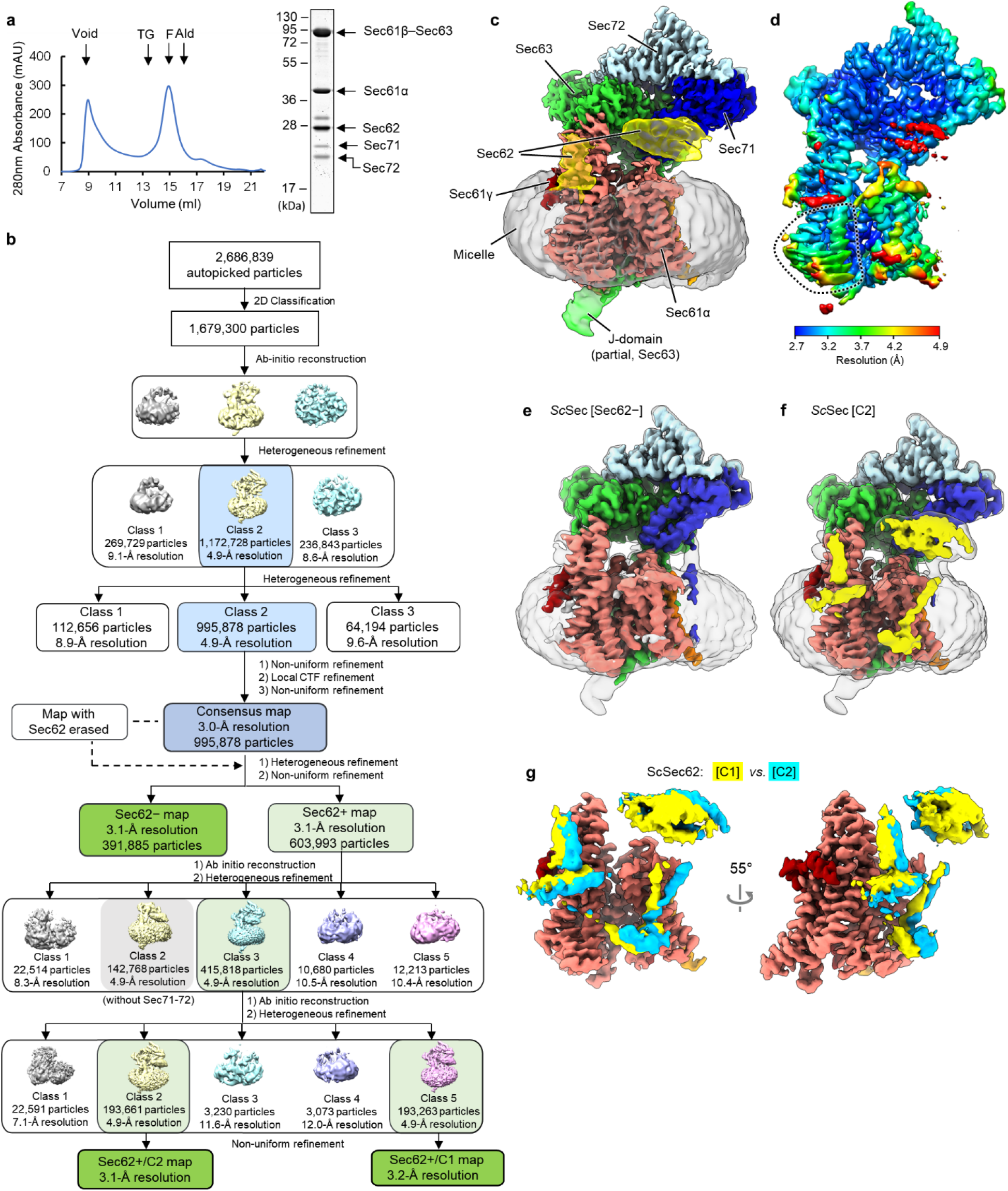
Cryo-EM analysis of the wild-type (WT) *S. cerevisiae* Sec complex (*Sc*Sec). **a**, Purification of the WT *Sc*Sec complex. Left, a chromatogram from Superose 6 size-exclusion chromatography of the affinity purified WT *Sc*Sec complex (MW standards: Tg, thyroglobulin; F, ferritin; Ald, aldolase). Right, the Coomassie-stained SDS-PAGE gel of the Superose 6 peak fraction. In this gel, Sec61γ (~10 kDa) migrated off the bottom. **b**, A diagram of the cryo-EM single particle analysis procedure. **c**, The 3.0-Å-resolution consensus map of *Sc*Sec. Salmon, Sec61α; orange, Sec61β; red, Sec61γ; yellow, Sec62; green, Sec63; blue, Sec71; light blue, Sec72; Grey, detergent micelle. Semitransparent surface, lowpass-filtered (5 Å for Sec62 and the J-domain and 7 Å for the micelle) maps shown at a lower contour level. **d**, As in c, but showing a local resolution map. Note that in addition to Sec62, the TM7-TM8 region of Sec61α (dotted line) displays noticeably lower solution than the overall resolution due to conformational heterogeneity (see Fig. 2a). **e**, As in c, but with the 3.1-Å-resolution map of the Sec62-class. Semitransparent surface, 6-Å-lowpass-filtered map at a lower contour level. **f**, As in c, but with the 3.1-Å-resolution map of the Sec62+/C2 class. **g**, The Sec62 densities of the C1 (yellow) and C2 (cyan) classes were compared after aligning the two cryo-EM maps. For simplicity, only Sec61 (from the C1 class) and Sec62 are shown.

**Extended Data Figure 2.**
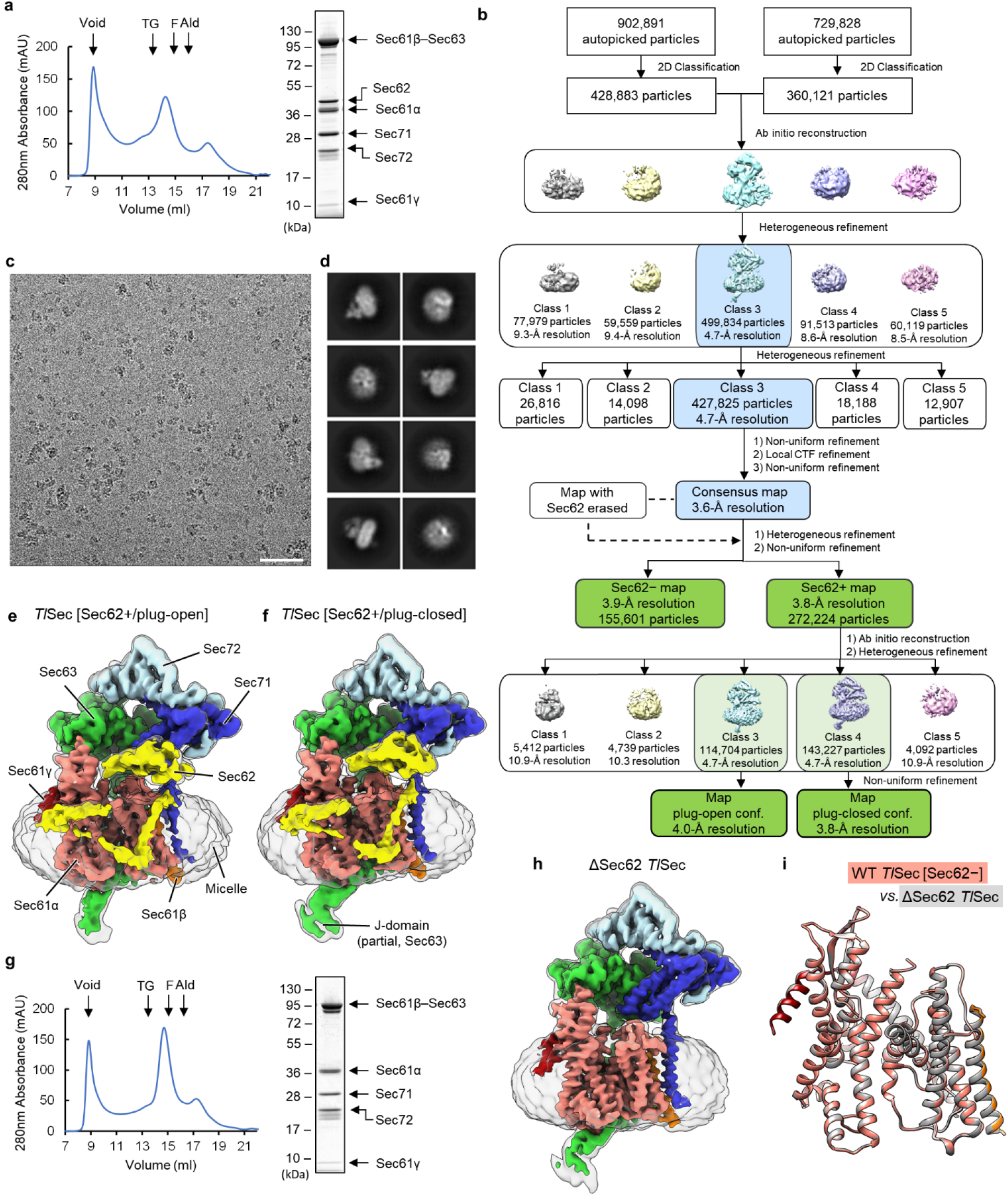
Cryo-EM analysis of the WT and ΔSec62 *T. lanuginosus* Sec complexes (*Tl*Sec). **a**, Purification of the WT *TlSec* complex. Left, a chromatogram from Superose 6 size-exclusion chromatography of the affinity purified WT *TlSec* complex. Right, the Coomassie-stained SDS-PAGE gel of the Superose 6 peak fraction. **b**, A diagram of the cryo-EM single particle analysis procedure. **c**, A representative cryo-EM micrograph. Scale bar, 50 nm. **d**, Examples of selected 2D class averages. **e**, The 4.0-Å-resolution map of the Sec62+/plug-open class of WT *TlSec*. The color scheme is the same as in Fig. 1. Semitransparent surface, a 7-Å-lowpass-filtered map shown at a lower contour level. **f**, As in e, but showing the 3.8-Å-resolution map of the Sec62+/plug-closed class. **g**, As in a, but purification of the ΔSec62 *TlSec* complex. **h**, As in e, but with the 3.7-Å-resolution map of ΔSec62 *TlSec* complex. **i**, The atomic models of the Sec61 complexes from the Sec62-class of WT *TlSec* (in color) and the ΔSec62 *TlSec* structure (in grey) were aligned and compared (RMSD of Cα atoms is 0.24Å).

**Extended Data Figure 3.**
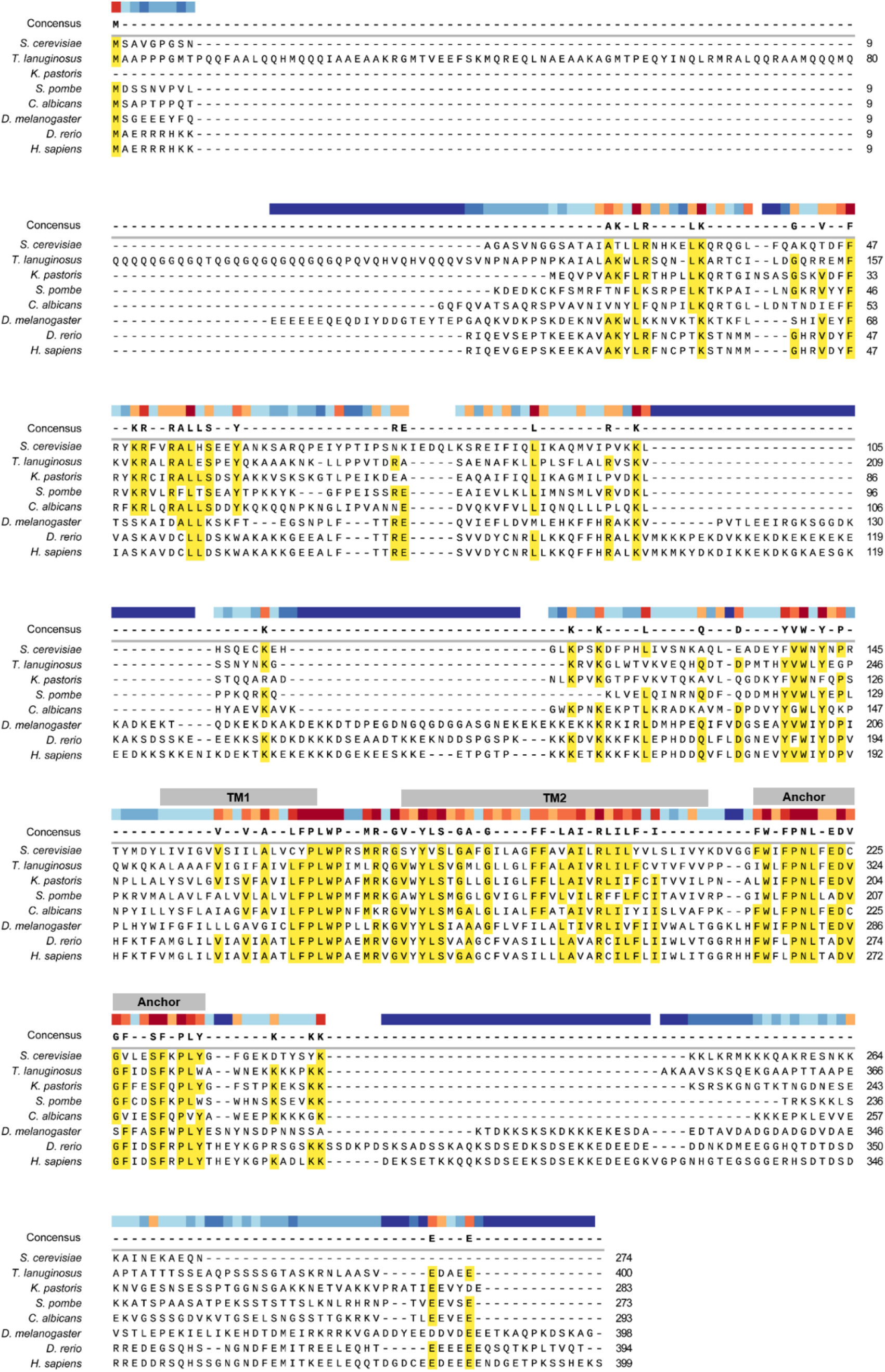
Sequence alignment of Sec62. Sec62 sequences from fungal and metazoan species were aligned. Approximate positions of the TM1, TM2, and anchor domain are indicated (grey). Color bar, heat map showing the degree of conservation (red, more conserved; blue, less conserved). Yellow highlight, amino acid positions with >50% sequence identity.

**Extended Data Figure 4.**
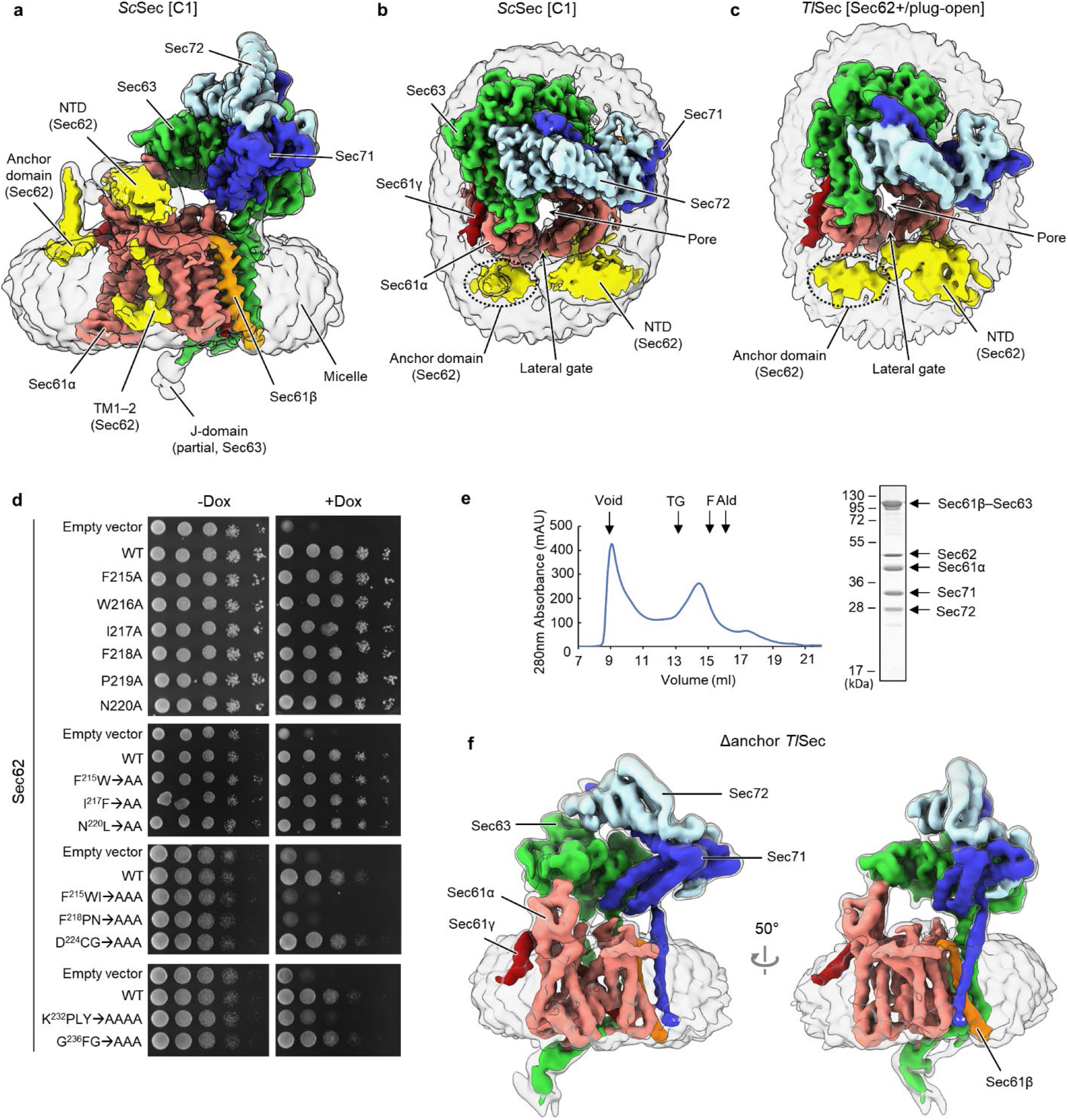
Structure and mutagenesis analysis of the anchor domain of Sec62. **a**, As in Fig. 1a, but a side view additionally showing a 6-Å-lowpass-filtered map at a lower contour level (semitransparent surface). **b**, As in a, but showing a view from cytosol. The anchor domain of Sec62 is indicated by a dotted oval. **c**, As in b, but showing the map of the Sec62+/plug-open class of *Tl*Sec. Semitransparent surface is a 7-Å-lowpass-filtered map. **d**, Yeast growth complementation tests for Sec62 anchor mutants. The yeast strain (ySI62) whose endogenous Sec62 is expressed under a tetracycline-repressible promoter was transformed with a CEN/ARS plasmid expressing WT or indicated mutant Sec62 under its native promoter. As a control, empty vector was used. In the right panels, 10 μg/mL doxycycline was included to repress the expression of endogenous Sec62. All growth assays were performed at 30°C. The top two panels (single and double mutants) were grown on synthetic complete (SC) medium lacking leucine, and the bottom two panels (triple and quadruple Ala mutants) were grown on YPD medium. **e**, As in Extended Data Fig 2a, but with the Δanchor mutant of *Tl*Sec. **f**, As in Extended Data Fig. 2e, but showing the 4.38-Å-resolution map of Δanchor *Tl*Sec. Semitransparent surface, 7-Å-lowpass-filtered map at a lower contour level. We note that the conformation of Δanchor *Tl*Sec is essentially identical to ΔSec62 *Tl*Sec.

**Extended Data Figure 5.**
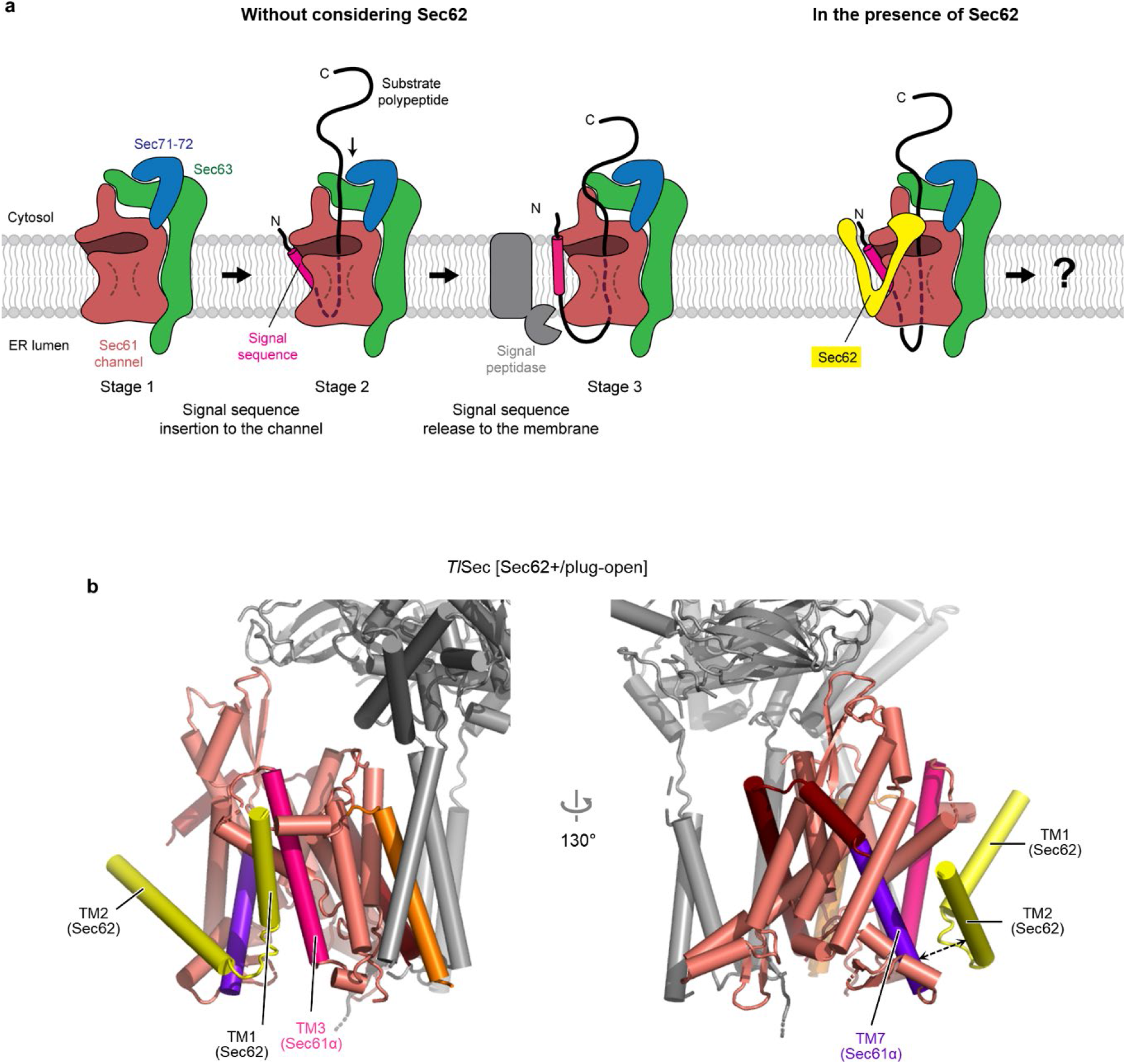
A potential path for signal sequence release from the lateral gate into the membrane. **a**, A schematic model for substrate insertion into the Sec complex and translocation. Substrates are expected to insert into the Sec61 channel as a loop with both N- and C-termini exposed in the cytosol. The N-terminal signal sequence may sit at the lateral gate (Stage 2) as seen in the structures of eukaryotic co-translational and bacterial post-translational translocation intermediates^10-12^. For later signal sequence cleavage by the signal peptidase, it is likely necessary for the signal sequence to be released into the membrane. In the presence of Sec62 (right), the signal sequence would become trapped at the lateral gate if Sec62 forms tight contacts with the lateral gate. **b**, As in Fig. 1e, but with the *Tl*Sec [Sec62+/plug-down] structure. Dashed arrow, a gap between TM2 of Sec62 and TM7 of Sec61α.

**Extended Data Figure 6.**
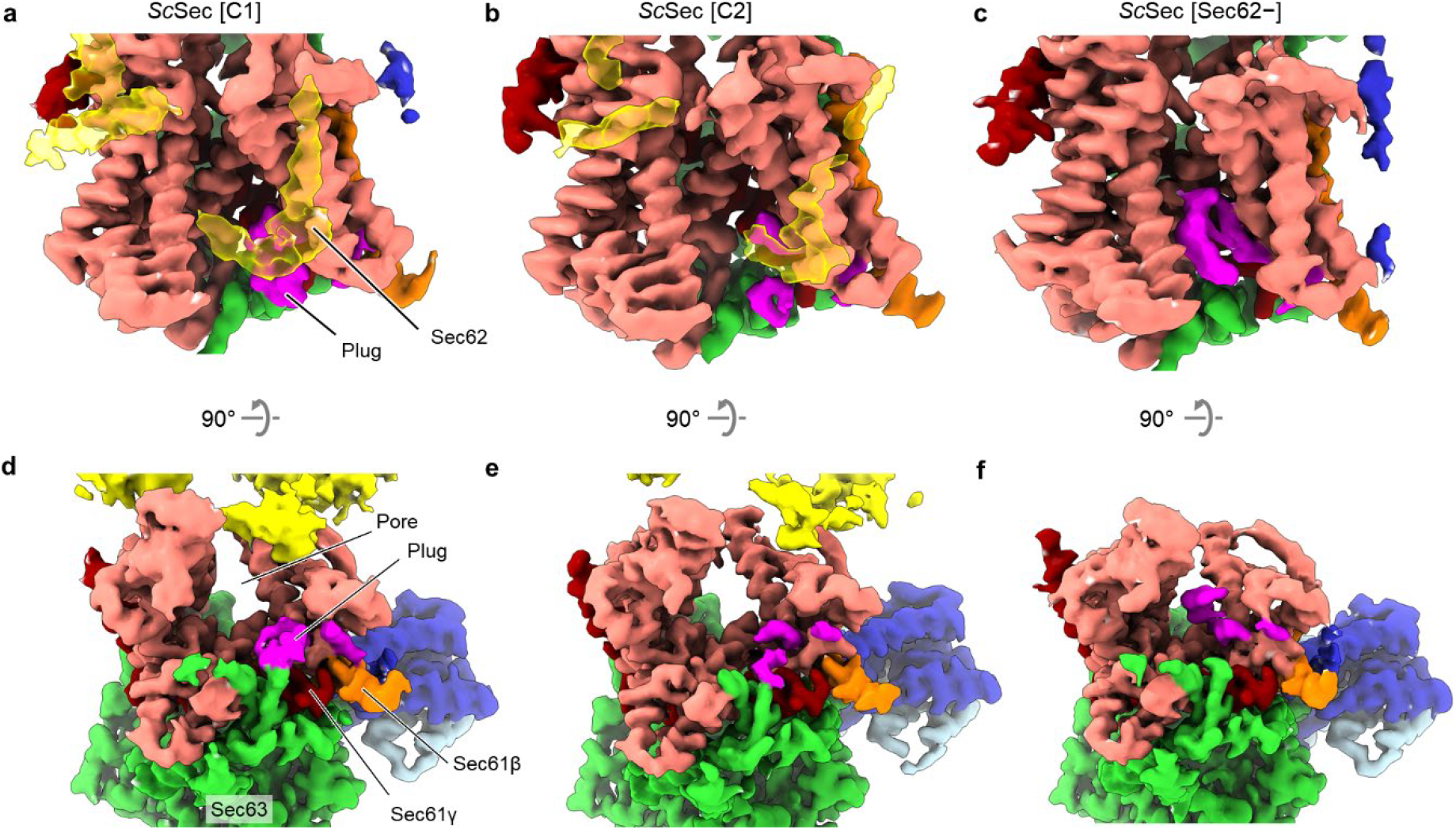
The presence of Sec62 induces opening of the vertical gate of Sec61 by displacing the plug domain from the closed position. **a-c**, Views into the lateral gate (front view) of the three classes of the WT *Sc*Sec structure, C1, C2, and Sec62-. Color scheme: salmon, Sec61α; orange, Sec61β; red, Sec61γ; yellow, Sec62; green, Sec63; blue, Sec71; light blue, Sec72; magenta, the plug domain. **d-f**, as in a-c, but showing views from the ER lumen.

**Extended Data Figure 7.**
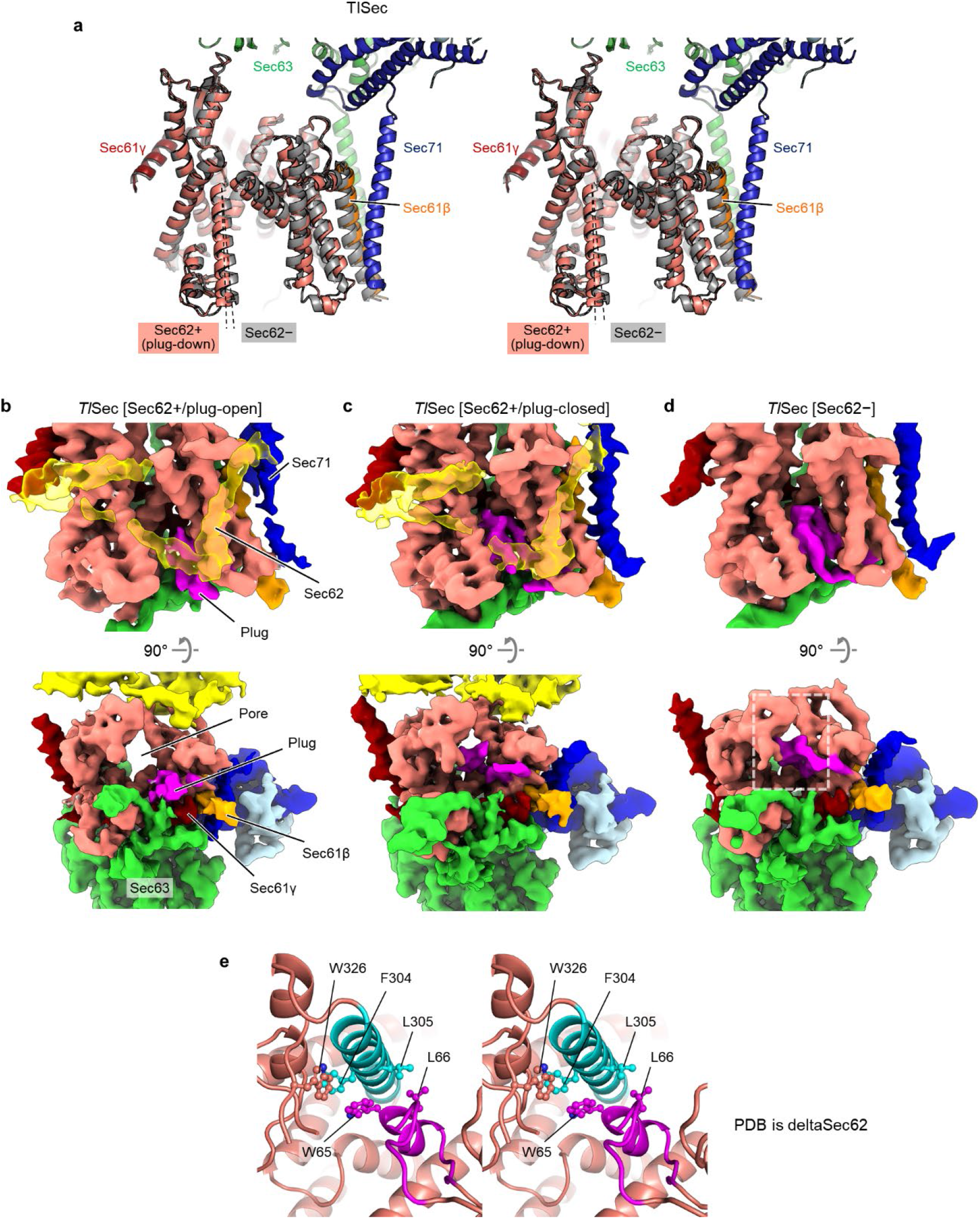
Conformational changes in the lateral gate and plug domain by Sec62 in the *TlSec* complex. **a**, As in Fig. 2a, but with the WT *Tl*Sec (stereo view). Two classes, Sec62+/plug-open (in color) and Sec62- (grey) are shown. Dashed line, TM7 of Sec61α. For simplicity, Sec62 and the plug are not shown. We note that in the Sec62+/plug-closed class, TM7 assumes an intermediate position of the plug-open and Sec62-classes. **b-d**, As in Extended Data Fig. 6 but with the WT *TlSec* structures. Shows are front views (upper panels) and views from the ER lumen (lower panels). We note that in both the plug-open and plug-closed classes, the conformation of Sec62 (yellow) is similar to that of the *Sc*Sec[C2] structure. The area indicated by a white dashed box in g are shown in e. **e**, A stereo view showing an interaction between the plug domain (magenta) and TM7 (cyan) of Sec61α. Side chains that are involved in the interaction are shown in a ball and stick representation. W326 belongs to the loop 7/8 of Sec61α. We note that a highly similar interaction is also present in the crystal structure of *P. furiosus* SecY (ref. ^36^; PDB ID 3MP7).

**Extended Data Figure 8.**
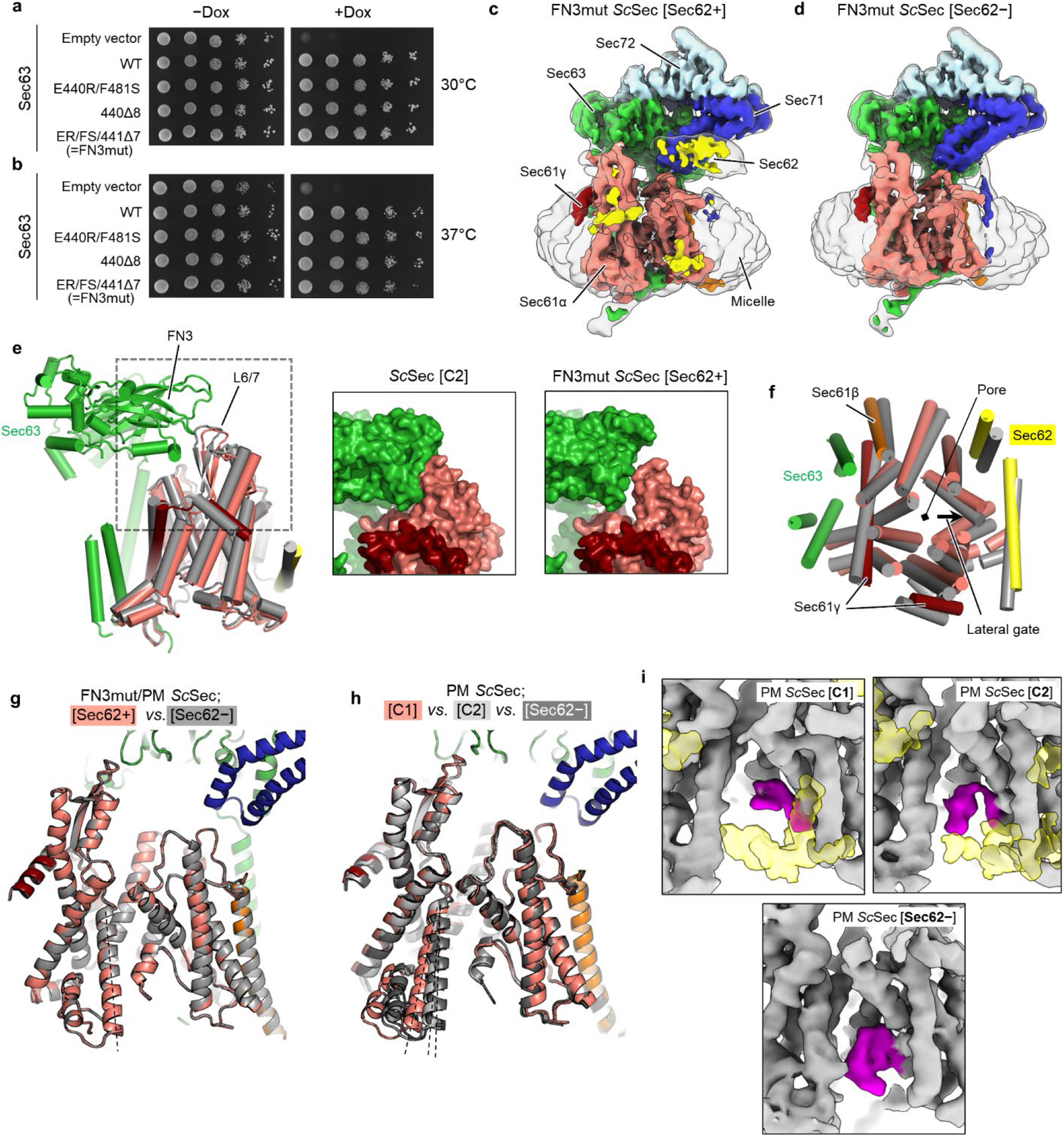
Structures of the *Sc*Sec complex containing mutations in the FN3 domain of Sec62 and the pore ring of Sec61α. **a**, The same yeast growth complementation experiment shown in Fig. 3b (left panel), but additionally showing a control without doxycycline. **b**, As in a, but the plates were incubated at 37°C. **c**, The 3.90-Å-resolution cryo-EM map of the Sec62+ class of the FN3mut *Sc*Sec complex. **d**, As in c, but with the 4.0-Å-resolution cryo-EM map of the Sec62-class of the FN3mut *Sc*Sec complex. **e**, The interaction between the FN3 domain of Sec63 and the L6/7 of Sec61α. Left, a side view showing Sec63 (green) and the Sec61 complex (grey, C2 class of WT *Sc*Sec; color, Sec62+ class of FN3mut *Sc*Sec). The structures were aligned with respect to Sec63. The area indicated by a grey dashed box is shown in the middle and right panels with a solvent-accessible surface representation. **f**, As in the left panel of e, but showing the cytosolic view into the Sec61 complex. **g**, As in Fig. 2a, but comparing the Sec62+ and Sec62-classes of FN3mut/PM *Sc*Sec. **h**, As in Fig. 2a, but comparing the three classes of PM *Sc*Sec. **i**, As in Fig. 3e, but with PM *Sc*Sec. We note that although the plug domain is partly visible in the C1 and C2 classes of PM *Sc*Sec, its density is substantially weaker than that of the Sec62-class.

**Extended Data Figure 9.**
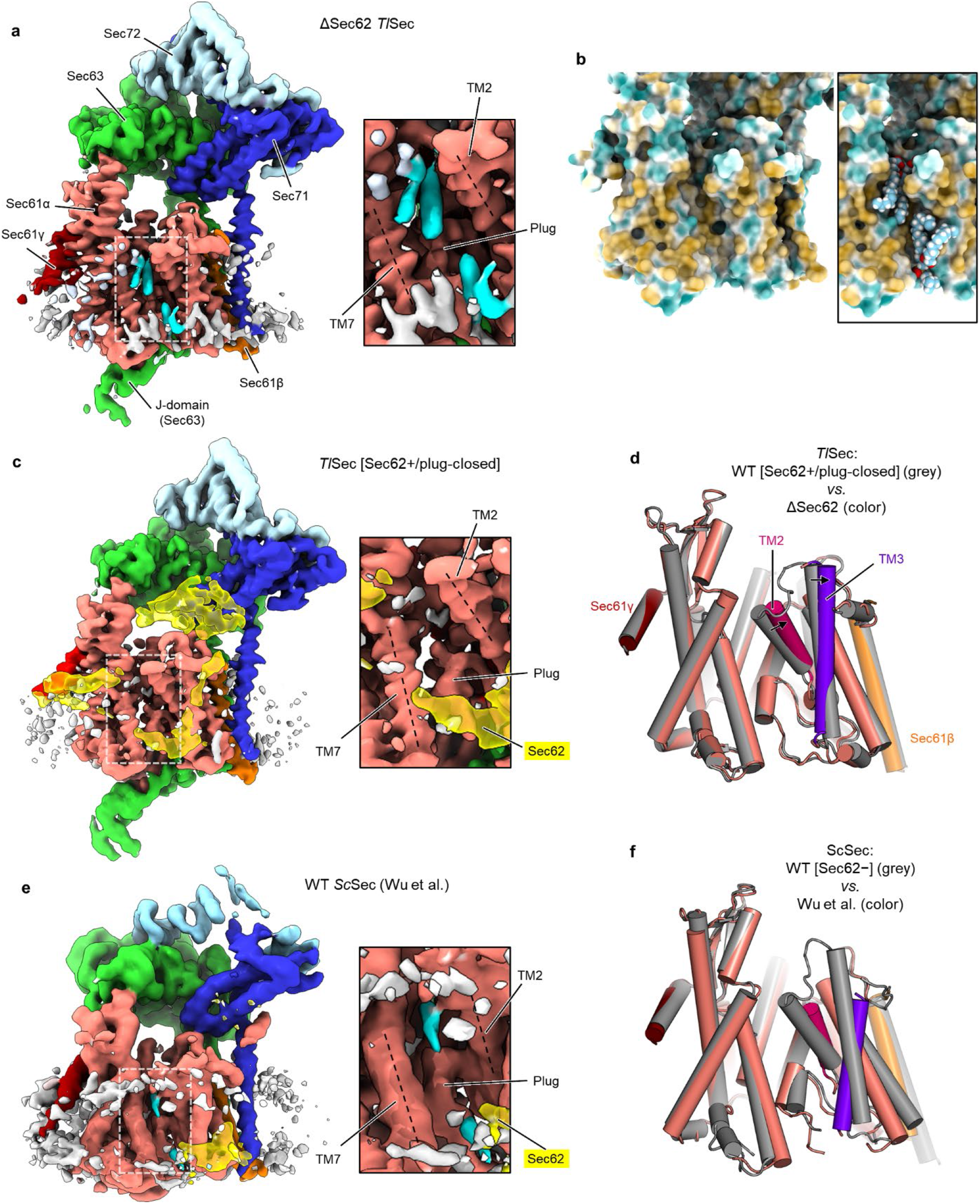
Lipid/detergent molecules at the lateral gate in the *Tl*Sec structure lacking Sec62 (ΔSec62). **a**, A view into the lateral gate of ΔSec62 *Tl*Sec. Left, a front view. Non-protein densities are in grey. Densities in cyan are lipid/detergent molecules intercalated at the lateral gate (an atomic model is shown in the right panel in b). Right, a zoomed-in view of the lateral gate (area indicated by the white dashed box in the left panel). **b**, A surface representation of the Sec61 complex (front view) showing the distribution of hydrophobic (yellow) and hydrophilic (cyan) amino acids. In the right panel, phosphatidylcholine lipid molecules modelled into the cryo-EM densities were additionally shown in a space-filling representation. **c**, As in a, but with the Sec62+/plug-closed class of WT *Tl*Sec. **d**, A comparison of the Sec61 atomic models of ΔSec62 *Tl*Sec (in color) and WT *Tl*Sec [Sec62+/plug-closed] (in grey). Movements of TM2 (purple) and TM3 (violet) are indicated. **e**. As in a, but the WT *Sc*Sec structure by Wu et al. (EMDB-0440; ref. ^29^). **f**, As in d, but comparing the structure by Wu et al. (in color; PDB 6ND1; ref. ^29^) and the *Sc*Sec[Sec62-] structure of the present study (in grey).

**Extended Data Figure 10.**
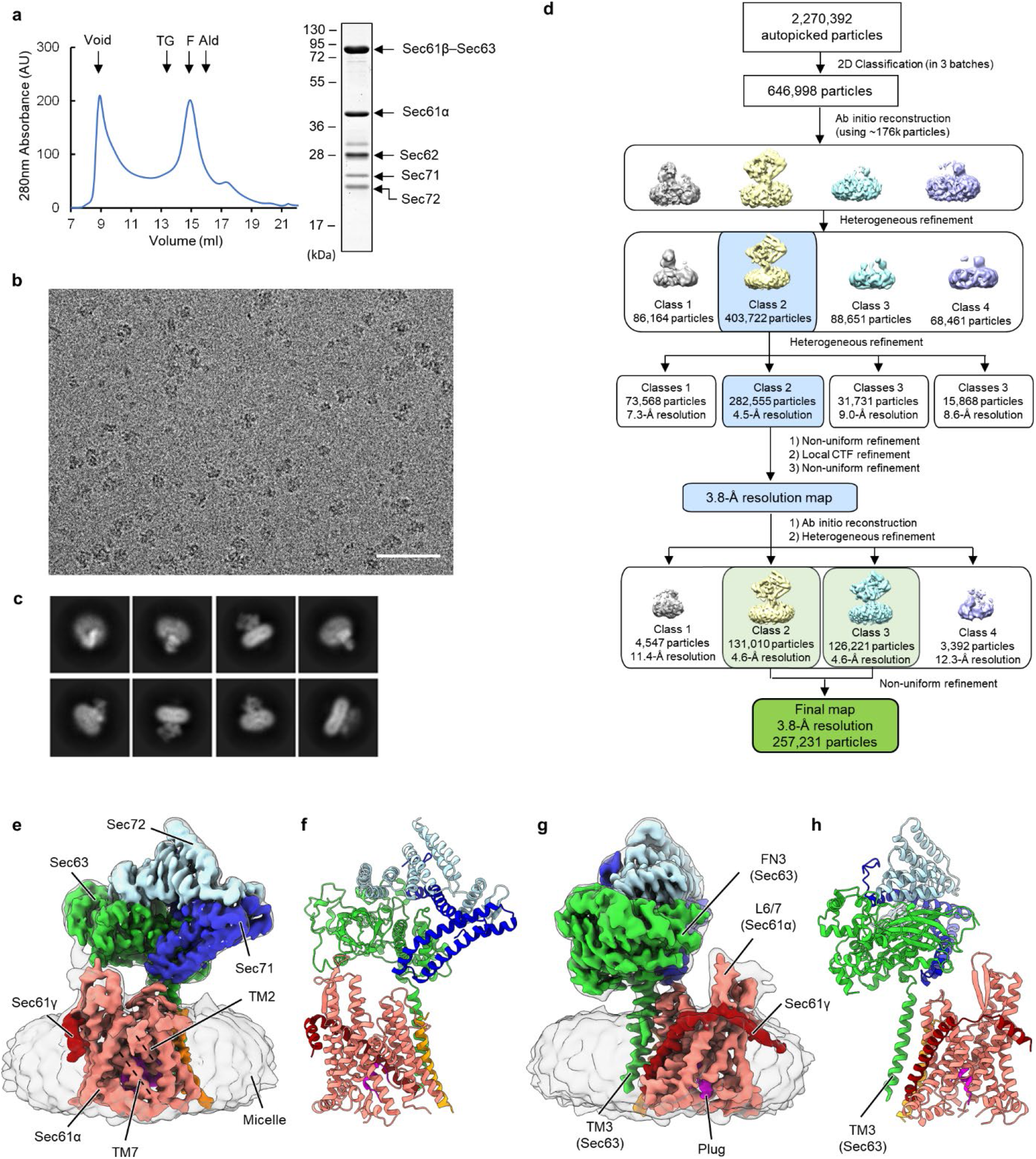
Structure of a fully closed *Sc*Sec complex containing FN3mut/Δ210-216 Sec63. **a**, Purification of the FN3mut/Δ210-216 *Sc*Sec complex. Left, a chromatogram from Superose 6 size-exclusion chromatography of the affinity purified mutant *Sc*Sec complex. Right, Coomassie-stained SDS-PAGE gel of the Superose 6 peak fraction. In this gel, Sec61γ (~10 kDa) migrated off the bottom. **b**, A representative cryo-EM micrograph. Scale bar, 50 nm. **c**, Examples of selected 2D class averages. **d**, A diagram of the cryoEM single particle analysis procedure. **e**-**h**, The cryo-EM map (e and g) and atomic model (f and h). The color scheme is the same as in Fig. 1 a,d. The plug domain is shown in magenta. Shown are front (e and f) and side (g and h) views.

## Methods

### Yeast strains

A list of yeast strains used in this study is given in Supplementary Table 1.

The yeast strain (ySI7) used for purification of the wildtype (WT) *Sc*Sec complex has been described previously^28^. Briefly, this strain expresses a fusion protein of Sec61β (Sbh1), Sec63, and a green fluorescent protein (GFP) from the genomic *SEC63* locus under the endogenous promoter of *SEC63* (the endogenous *SBH1* copy is deleted). The C-terminus of Sbh1p and the N-terminus of Sec63 are separated by a 15-amino-acid-long glycine-serine linker. There is a flexible linker containing a Tobacco etch virus (TEV) cleavage sequence between the C-terminus of Sec63 and GFP.

To enable purification of the PM (pore mutant) *Sc*Sec complex, we generated strain ySI8 by modifying ySI7. We first clone the *SEC61* gene (from 1965-bp upstream to 668-bp downstream of the Sec61 coding sequence (CDS) of BY4741) into a pBlueScript-derived cloning vector (comprised of a pUC origin and an ampicillin resistance gene). We then inserted a *LEU2* selection marker cassette (amplified by PCR using pYTK075 as a template, a forward primer: 5’-agctaaataagatctTCGAGGAGAACTTCTAGTATATCTACATAC, and a reverse primer: 5’-tatatataggagctcCTGCCTATTTAACGCCAAC; uppercase for *LEU2*-specific sequences and lowercase for *SEC61*-specific sequences) was inserted between 125 bp and 126 bp downstream of the Sec61 stop codon by In-Fusion cloning (Takara Bio). Pore mutations (M90L/T185I/M294I/M450L) were introduced to Sec61 by site-specific mutagenesis. The resulting plasmid was then linearized by cutting the plasmid backbone with *NotI.* The DNA fragment was introduced into ySI7 by a standard lithium acetate/polyethylene glycol transformation protocol. Recombinants were selected on a leucine drop-out synthetic complete [SC(-Leu)] agar medium. Incorporation of the mutations was verified by PCR and Sanger sequencing of single colonies.

To purify the FN3mut *Sc*Sec complex, we generated strain ySI73 by modifying strain TH_5187 (Horizon Discovery) from the Hughes collection^45^, where the expression of chromosomal Sec63 is under the control of a tetracycline repressible promoter. First, the endogenous copy of *SBH1* of TH_5187 was replaced with a hygromycin resistance marker (hphMX) cassette. An hphMX cassette fragment was amplified by PCR using pFA6a-hphMX6 (ref. ^46^) as a template (forward primer: 5’-gggaaaagatttcaaccaccacttcaaaacaccacactctacctcctaccatactccataAGCTTGCCTCGTCCCC; reverse primer: 5’-tagtcttgttttgtcaaatagggtggataaaagctgaatcattactgaagaaaattcttaCAGTATAGCGACCAGCATTCAC; uppercase for vector specific sequences and lowercase for sequences homologous to yeast chromosomal sequences). The DNA fragment was transformed into TH_5187. Single colonies were isolated from YPD (1% yeast extract, 2% peptone, 2% glucose) agar plates containing 400 μg/ml hygromycin (Gold Biotechnology), and integration was verified by PCR. The resulting strain (ySI48) was then further modified by integration of the FN3mut Sec63 construct into the *HO* locus using the transforming plasmid pSI74 (see below) linearized with *HindIII*, which cuts at the *E. coli* kanamycin resistance marker. Transformed cells were selected on SC(-Leu) agar medium, and integration was verified by PCR.

For purification of the FN3mut/PM *Sc*Sec complex, we used strain ySI74. To generate ySI74, we first modified TH_5187 to contain the pore mutations in the *SEC61* gene, similarly as described above for ySI8, but using a nourseothricin resistance cassette (natMX6) instead of the *LEU2* marker. The natMX6 cassette was amplified from pFA6a-natMX6 (ref. ^46^) and inserted into the pBlueScript-Sec61 plasmid at 125 bp downstream of the Sec61 stop codon. After transformation of the linearized plasmid, recombinants were selected on YPD agar plates containing 100 μg/ml nourseothricin (Gold Biotechnology), resulting in ySI42. Subsequently, *SBH1* deletion and Sec63 FN3mut mutation were introduced to the strain as described for ySI73.

For purification of the FN3mut/Δ210-216 *Sc*Sec complex, we used strain ySI112. To generate ySI112 we modified strain ySI48 (TH_5187 *sbhlΔ::hphMX6*) by integration of the FM3mut/Δ210-216 Sec63 construct into the *HO* locus using the transforming plasmid pSI120 (see below) linearized with *Hind*III.

The WT and mutant *Tl*Sec complexes were expressed using the yeast strains ySI67 (for WT), ySI77 (for ΔSec62), and ySI113 (for Δanchor). These strains were generated from the parental strain yMLT62 (a gift from J. Thorner; ref. ^47^), which expresses the β-estradiol-responsive chimeric transcription activator Gal4dbd.ER.VP16. All *Tl*Sec subunits were co-expressed with an integration vector (pYTK-e101) generated using MoClo Yeast ToolKit (YTK)^48^ (see below). Expression of each gene is driven by a GAL1 promoter. pYTK-e101 contains a nourseothricin resistance marker and *URA3* homology sequences for chromosomal integration. pYTK-e101 encoding WT (pSI65) or mutant *Tl*Sec (pSI87 for ΔSec62 and pSI94 for Δanchor) were linearized with *NotI* and transformed into yMLT62. Recombinants were selected by growth in YPD agar plates containing 100 μg/ml nourseothricin. Integration was verified by PCR as described^48^.

For yeast growth complementation assays for Sec62, we generated ySI62, whose chromosomal Sec62 expression can be repressed in the presence of doxycycline. The tetracycline response element (TRE) as well as the upstream kanMX cassette were PCR amplified from genomic DNA of TH_5187 with primers containing 60 bps overhangs homologous to the N-terminus of Sec62 (forward primer: 5’-gacggaatagacgtgtcgttttcccaatactggcatacaaatcaagagggagaagagtggGGCGTTAGTATCGAATCG; reverse primer: 5’ - tgtagcagatccgccattgacactagcacctgcattgctacctggacctacggctgacatGGATCCCCCGAATTG; uppercase for sequence specific to TRE-kanMX and lowercase for sequences homologous to yeast chromosomal sequences). The amplicon was then transformed into the strain R1158 (the parental strain for TH_5187) which contains the “tet activator” (tTA). Transformed cells were selected on 300 μg/ml G418 (Fisher Chemical) containing YPD-agar plates and integration was verified by PCR.

Strain ySI89 was used for complementation assays testing synthetic growth defects of FN3mut Sec63 and Sec61 mutants (Fig. 3c). To generate this strain, TH_4087 (Hughes strain with the chromosomal *SEC61* expressing under a tetracycline-repressible promoter) was modified such that its endogenous Sec63 was mutated to FN3mut. A DNA segment encoding part of FN3mut Sec63 was amplified by PCR from plasmid pSI16 (from 164th amino acid to the stop codon) and inserted to pFA6a-natMX6 immediately before an ADH1 terminator, which precedes the natMX6 cassette. The resulting construct was amplified by PCR to include 773 bp upstream of the first mutated amino acid (E440R) and 50 bp of the 3’ untranslated region of the *SEC63* locus (forward primer: 5’ - CCCTTACTGACGAATTGGTTAGGC; reverse primer: 5’-atgtatctatttttataaagatgaaatatatacgtctaagagctaaaatgGGCCGCATAGGCCACTAG; uppercase for sequence specific to the plasmid and lowercase for sequences homologous to yeast chromosomal sequence). The amplicon was transformed into TH_4087 and selected on 100 μg/ml nourseothricin containing YPD-agar plates. Incorporation of the mutation was verified using Sanger sequencing.

### Plasmids

A list of plasmids used in this study is given in Supplementary Table 1.

The integration vectors pYTK-e101 and pYTK-e106, and CEN/ARS plasmid pYTK-e112 were generated using Golden Gate *BsaI* assembly of parts from MoClo YTK^48^ (for pTYK-e101, parts were 8, 47, 73, 78, 86, 90, and 92; for pYTK-e106, part numbers were 8, 47, 73, 75, 88, 90, and 94; for pYTK-e112, part numbers were 8, 47, 73, 75, 81, and 84). pYTK-e101 contains a natMX6 marker and integrates into the *URA3* locus. pYTK-e106 contains a *LEU2* auxotroph marker and integrates into the *HO* locus. pYTK-e112 contains a *LEU2* auxotroph marker.

The *Tl*Sec-expressing pYTK-e101 plasmids were generated using MoClo YTK as follows. First, gene fragments encoding *Tl*Sec subunits were chemically synthesized based on protein sequences of *T. lanuginosus* ATCC 2000065 (https://gb.fungalgenomics.ca) and cloned into the YTK entry vector pYTK001 (ref. ^48^). Codons were optimized for yeast. In the case of the Sec63 and Sec61β subunits, a fusion construct (*Tl*Sec61β-GGSGGSGGSGGSGGS-*Tl*Sec63-TEV-GFP) was synthesized similarly to the expression of the *Sc*Sec complex. Each synthesized CDS was then cloned into the pYTK095 vector (ref. ^48^) as an expression cassette together with a *GAL1* inducible promoter, an *ENO1* terminator, and connector parts by Golden Gate *BsaI* assembly. Subsequently, the multigene expression construct was generated by Golden Gate *BsmBI* assembly of the pYTK095 plasmids and pYTK-e101 resulting in pSI65 (the Sec gene placed in tandem in the following order: *Tl*Sec61α, *Tl*Sec61γ, *Tl*Sec62, *Tl*Sec61β-*Tl*Sec63-GFP, *Tl*Sec71, and *Tl*Sec72). For ΔSec62 *Tl*Sec (plasmid pSI87), pYTK095-*Tl*Sec62 was replaced by a non-expressing spacer cassette in the *BsmBI* assembly. For Δanchor *Tl*Sec (plasmid pSI94), amino acid residues N^319^LF…WNE^338^ of *Tl*Sec62 were replaced with a Gly/Ser linker (GGSGGSGGS) before the multigene *Bsm*BI assembly.

For expression of Sec63-mutant *Sc*Sec complexes (FN3mut and FN3mut/Δ210-216), WT *Sc*Sec63 was first amplified from genomic DNA of BY4741 by PCR to include the endogenous promoter and terminator (187 bp upstream and 97 downstream of the CDS) and cloned into pYTK-e112 between the two *Bsa*I sites (pSI5). This plasmid was then further modified to have Sbh1 and GFP flanking Sec63 as in other Sbh1-Sec63 fusion constructs (i.e., Sbh1–GGSGGSGGSGGSGGS–Sec63–TEV–GFP). Then FN3mut (E440R/F481S/Δ441-447) or FN3mut/Δ210-216 (FN3mut and residues L^210^PRFLVD^216^, replaced with SGSGGSG) mutations were introduced by site-specific mutagenesis. For generation of strains ySI74 and ySI112 (chromosomal integration of mutant Sec63 to the HO locus), the expression cassette for Sbh1-Sec63-GFP was transferred to pYTK-e106 by restriction digestion and ligation.

For growth complementation assays, *SEC61* (710 bp upstream to 264 bp downstream of CDS) and *SEC62* (251 bp upstream to 123 bp downstream of CDS) were amplified by PCR using genomic DNA of BY4741 and cloned into pYTK-e112, resulting in pSI123 and pSI39, respectively. Plasmids used for growth complementation assays of Sec63 mutants were derived from pSI5 (see above) by adding a TEV-GFP tag at the C-terminus of Sec63 (the Sbh1 fusion was not introduced, and thus the constructs have the native N-terminus).

### Yeast growth complementation assays

The yeast strains were transformed with each pYTK-e112 plasmid encoding the indicated protein under its endogenous promoter. Cells were selected on SC(-Leu) agar medium. Single colonies were picked and grown overnight in SC(-Leu) medium. The cultures were diluted with water to OD600 of 1.0 and further serially diluted by factors of 10 (in Fig 3b,c, starting concentration was OD600 of 0.1). The diluted cultures (10 μL each) were spotted on SC(-Leu) agar plates. In the case of Extended Data Fig. 4 (bottom two panels only), YPD agar medium was used. Where indicated, plates contained 10 μg/ml doxycycline. Plates were incubated at 30°C unless otherwise stated. The following strains were used for the indicated experiment: TH_5187 (Fig. 3b left panel, Fig. 4, and Extended Data Fig. 8a,b), ySI42 (Fig. 3b right panel), ySI89 (Fig. 3c), and ySI62 (Extended Data Fig. 4c)

### Protein purifications

For purification of *Sc*Sec complexes, yeast cells (ySI7 for WT, ySI8 for PM, ySI73 for FN3mut, ySI74 for FN3/Δ210-216) were grown in YPD medium to OD600 of 2-3, before harvest. For purification of *Tl*Sec, cells were grown in YPD medium to OD of 1.0. After adding 50 nM β-estradiol, cells were further grown until reaching OD 2-3. All cultures were grown in 30°C, except for the FN3mut/PM and FN3/Δ210-216 variants of *Sc*Sec, for which cells were grown at 22°C. Cells were harvested by centrifugation (8 min at 6,400 xg), washed once with ice-cold tris-buffered saline (20mM Tris pH 7.5 and 150 mM NaCl), frozen in liquid nitrogen, and stored at −75°C before use.

All *Sc*Sec and *Tl*Sec complexes were purification as described previously^28^. Briefly, cells were lysed by cryo-milling and resuspended in buffer containing 50 mM Tris pH 7.5, 200 mM NaCl, 1 mM EDTA, 10% glycerol, 2mM DTT, 5 μg/ml aprotinin, 5 μg/ml leupeptin, 1 μg/ml pepstatin A, and 1.2 mM PMSF. Membranes were solubilized by adding 1% lauryl maltose neopentyl glycol (LMNG; Anatrace) and 0.2% cholesteryl hemisuccinate (CHS; Anatrace) directly to the whole-cell lysate for 1.5 h at 4°C. The lysate was clarified by ultracentrifugation at 125,000 xg for 1 h. The Sec complex was bound to agarose beads conjugated with anti-GFP nanobody and the buffer was exchanged to 50 mM Tris pH 7.5, 200mM NaCl, 1.0 mM EDTA, 2 mM DTT, 0.02% glycol-diosgenin (GDN; Anatrace), and 10% glycerol. The complex was eluted by incubating the beads with TEV protease (~10 μg/ml) overnight and further purified by size-exclusion chromatography (Superose 6 Increase; GE Lifesciences) in 20 mM Tris pH 7.5, 100 mM NaCl, 1mM EDTA, 2 mM DTT, and 0.02% GDN. Peak fractions were concentrated to ~5 mg/ml and used immediate for cryo-EM. We note that the yields of all mutant *Sc*Sec complexes were comparable to that of the WT complex.

### Cryo-EM grid preparation and data collection

Purified samples were supplemented with 3 mM fluorinated Fos-Choline-8 (FFC8; Anatrace) before plunge freezing. The samples were applied on holey carbon gold grids (Quantifoil 1.2/1.3, 400 mesh) that were glow discharged for 35 seconds using PELCO easiGlow glow discharge cleaner. Plunge freezing was performed using Vitrobot Mark IV (FEI) set at 4°C and 100% humidity. Whatman No. 1 filter paper was used to blot the samples.

Datasets for *Tl*Sec, ΔSec62 *Tl*Sec and FN3mut/Δ210-216 *Sc*Sec were collected on an FEI Talos Arctica electron microscope operated at an acceleration voltage of 200 kV. Datasets for WT *Sc*Sec, PM *Sc*Sec, FN3mut *Sc*Sec, FN3mut/PM *Sc*Sec and Δanchor *Tl*Sec were collected on an FEI Titan Krios electron microscope operated at an acceleration voltage of 300 kV and equipped with a Gatan Quantum Image Filter (a slit width of 20 eV). Both microscopes operated using SerialEM software^49^. Movies were recorded on a Gatan K3 Summit direct electron detector under the super-resolution mode (with a physical pixel size of 1.14 Å for *Tl*Sec and ΔSec62 *Tl*Sec, 0.9 Å for FN3mut/Δ210-216 *Sc*Sec and 1.19 Å for WT *Sc*Sec, FN3mut *Sc*Sec, FN3mut/PM *Sc*Sec and Δanchor *Tl*Sec,) with the exception of PM ScSec, which utilized a Gatan K2 Summit direct election detector (with physical pixel size of 1.15 Å). The samples were exposed to a total dose of ~50 e^-^ per Å^2^ applied over 42 frames. Defocus target was typically set between −0.8 μm and −2.4 μm. For detailed parameters, also see Tables 1 and 2.

### Cryo-EM image analysis

Micrographs collected from the microscopes were preprocessed by Warp^50^. Movie stacks were corrected for gains and subjected to tile-based motion correction and CTF estimation (7 by 5 tiles for datasets from the K3 detector and 5 by 5 for datasets from the K2 detector). Particles were automatically picked using the BoxNet algorithm of Warp. Low quality micrographs and particles, such as those containing crystalline ice or showing excessive motion blur, were removed by manual inspection. Motion corrected movies were exported with 2x-pixel- and 2x-frame-binning. Local particle motion corrections were performed in cryoSPARC v2 (ref. ^51^) after importing particle metadata and motion-corrected movie stacks. Box sizes of extracted particle images were 256 pixels except for the FN3mut/Δ210-216 *Sc*Sec dataset, which was 320 pixels. All subsequent single-particle analyses were performed with cryoSPARC v2 as described below. In the cases of the WT and FN3mut/Δ210-216 *Sc*Sec datasets, particle images extracted from Warp were directly used without local motion correction.

#### (1) WT *Sc*Sec

The single-particle analysis procedure for WT *Sc*Sec is outlined in Extended Data Fig. 1b. First, 2,686,839 picked particles were subjected to 2D classification, where empty micelles and classes of poor quality were removed. Selected 1,679,300 particles were then subjected to ab-initio reconstruction to yield three initial models, followed by heterogenous refinement using the initial maps (unless state otherwise, particle images were 2x scaled down to 128 by 128 pixels in all heterogenous refinement). Features of the Sec complex appeared in only one class (1,172,728 particles), particles of which were subjected to a second iteration of heterogenous refinement with the three classes from the first heterogenous refinement as references to further remove poor-quality particles. The resulting 995,878 particles were then subjected to a round of non-uniform refinement, local CTF refinement, and another round of non-uniform refinement, yielding a map at 2.98-Å resolution (consensus map). In order to separate the particles into classes containing and lacking Sec62, the N-terminal cytosolic domain (NTD) density of Sec62 in the consensus map was manually erased used UCSF Chimera^52^ and was used alongside the consensus map as initial references for heterogenous refinement. This yielded two classes: *Sc*Sec[Sec62-] with 391,885 particles which is largely devoid of detectable Sec62 and *Sc*Sec[Sec62+] with 603,993 particles. After non-uniform refinement, both classes refined to resolution of 3.07 Å. To further separate into subclasses containing different conformations of Sec62, the particles of *Sc*Sec[Sec62+] were subjected to a round of ab-initio reconstruction and heterogeneous refinement to yield five new classes. This step produced two major classes: one lacking Sec71-Sec72 (142,768 particles), and one showing the full complex features (415,818 particles). Particles of the latter class were subjected to a second round of ab-initio reconstruction and heterogenous refinement, yielding five new classes. Of these, two major classes showing the prominent features of the Sec complex (the other three classes did not show clear features of *Sc*Sec) were further refined using non-uniform refinement to yield the final maps of *Sc*Sec[C1] (from 193,263 particles) and *Sc*Sec[C2] (from 193,661 particles) at overall resolutions of 3.16 and 3.14 Å, respectively.

#### (2) PM *Sc*Sec

The PM ScSec dataset was analyzed using essentially the same procedure as for WT *Sc*Sec but starting with a dataset of 195,915 auto-picked particles (see Supplementary Fig. 1). After a round of 2D classification, ab-initio reconstruction, and heterogeneous refinement, the consensus class (91,813 particles) was obtained, which was subjected to non-uniform refinement to yield a 3.53-Å-resolution map. As with WT *Sc*Sec, the particles were further classified to [Sec62-] and [Sec62+] classes by heterogeneous refinement (35,573 and 56,240 particles, respectively), and the structures were refined to maps at resolutions of 4.02 and 3.78 Å, respectively. Particles of the [Sec62+] class was further classified by ab-initio reconstruction and heterogeneous refinement (five classes). One class (13,752 particles) lacked the Sec71-72 subunits, and the main class (36,506 particles) showed features of the full complex. The particles from the latter class were subjected another round of ab-initio reconstruction and heterogenous refinement, yielding two main classes, PM *Sc*Sec[C1] (17,341 particles) and PM *Sc*Sec[C2] (16,679 particles), which were further refined with non-uniform refinement to overall resolutions of 4.06 and 4.04 Å, respectively.

#### (3) FN3mut *Sc*Sec

The single-particle analysis procedure is outlined in Supplementary Fig. 1). The initial set of 1,274,219 auto-picked particles were subjected to 2D classification. After discarding empty micelle classes and classes showing poor features (resulting in 412,129 particles), we generated five initial models with ab-initio reconstruction. Only one class showed features of the Sec complex. Particles were subjected to two rounds of heterogeneous refinement to further remove particles of poor quality. The resulting 202,091 particles used for non-uniform refinement, which was followed by local CTF refinement and a second round of non-uniform refinement to obtain a consensus map at 3.73-Å resolution. Like with WT *Sc*Sec, these particles were further classified into two classes, one with Sec62 (FN3mut *Sc*Sec[Sec62+], 119,420 particles) and the other without Sec62 (FN3mut *Sc*Sec[Sec62-], 82,671 particles) using the consensus map and Sec62-NTD-erased map as initial references for heterogeneous refinement. FN3mut *Sc*Sec [Sec62+] and [Sec62-] particles were separately subjected to local CTF refinement and non-uniform refinement to yield final maps at 3.90- and 4.01-Å resolution, respectively. Further 3D classification of particles from the [Sec62+] class did not result in classes with a significant conformational difference.

#### (4) FN3mut/PM *Sc*Sec

The single-particle analysis procedure is outlined in Supplementary Fig. 1). The analysis was processed similarly as FN3mut *Sc*Sec. The initial set of 267,541 auto-picked particles were first cleaned up by 2D classification. The resulting 146,399 particles were subjected to ab-initio reconstruction (three classes). Only one main class showed features of the Sec complex. The 146,399 particles were then subjected to two rounds of heterogeneous refinement to remove non-Sec-complex particles. The resulting 86,843 particles were then subjected to non-uniform refinement, which was followed by local CTF refinement and a second round of non-uniform refinement to obtain a consensus map at 3.73-Å resolution. The particles were further classified to [Sec62+] and [Sec62-] classes (54,139 and 32,704 particles, respectively) with heterogeneous refinement, and final maps of PM/FN3mut *Sc*Sec [Sec62+] and [Sec62-] at 3.99- and 4.35-Å resolution respectively were obtained by non-uniform refinement followed by local CTF refinement and a second round of non-uniform refinement. Further 3D classification of particles from the [Sec62+] class did not yield classes with a significant conformational difference.

#### (5) FN3mut/Δ210-216 *Sc*Sec

The single-particle analysis procedure is outlined in Extended Data Fig. 10d. The initial set of 2,270,392 auto-picked particles were cleaned up by 2D classification. The resulting 646,998 particles were used to generate four initial maps with ab-initio reconstruction. Only one main class showed features of the Sec complex. Two rounds of heterogeneous refinement (with particle image 2x scaled down to 160 by 160 pixels) were performed to enrich particles of the Sec complex. The resulting 282,555 particles were subjected to non-uniform refinement, followed by local CTF refinement and a second round of non-uniform refinement to produce a 3.80-Å resolution map. The particles were then subjected to a second round of ab-initio reconstruction and heterogeneous refinement to generate four classes. Out of these classes, two showed features of the Sec complex (131,010 and 126,221 particles), maps of which were nearly identical. Particles of the two classes were combined for non-uniform refinement to yield the final map at 3.75-Å resolution.

#### (6) WT *Tl*Sec

The single-particle analysis procedure is outlined in Extended Data Fig. 2b. The initial set of 1,632,719 auto-picked particles were subjected to 2D classification in two batches to remove empty micelles and poor-quality particles. The resulting 789,004 particles were used to generate five initial 3D maps with ab-initio reconstruction. Only one (main) class showed features of the Sec complex. The 789,004 particles were subjected to heterogeneous refinement using the initial maps as references, which was followed by a second round of heterogeneous refinement. The resulting main class (427,835 particles) was refined using non-uniform refinement, local CTF refinement and second non-uniform refinement yielding a consensus map at 3.61-Å resolution. As with WT *Sc*Sec, particles were further classified to [Sec62+] and [Sec62-] classes with heterogeneous refinement using the consensus map and a Sec62-NTD-erased map as references (272,224 and 155,601 particles, respectively). The classes were further refined with non-uniform refinement yielding a 3.88-Å-resolution map of *Tl*Sec[Sec62-] and a 3.75-Å-resolution map of *Tl*Sec[Sec62+]. Particles of the [Sec62+] class (272,224 particles) were further subjected to ab-initio reconstruction and heterogeneous refinement (five classes). Two major classes (114,704 and 143,227 particles) showed the features of the Sec complex, which were further refined to the final maps of [Sec62+/plug-open] and [Sec62+/plug-closed] at overall resolutions of 4.02 and 3.76 Å, respectively. Unlike WT *Sc*Sec, a class lacking Sec71-Sec72 was not identified.

#### (7) ΔSec62 *Tl*Sec

The single-particle analysis procedure is outlined in Supplementary Fig. 1. The initial set of 546,712 auto-picked particles was subjected to two rounds of 2D classification with removal of poor classes in each round, resulting in 258,743 particles. Five initial 3D models were generated from the 258,743 selected particles by ab-initio reconstruction and further refined by heterogenous refinement. This produced two major classes (77,524 and 114,523 particles) which showed features of the Sec complex. Particles from the two classes were combined and refined with non-uniform refinement, local CTF refinement, and second non-uniform refinement, yielding the final map at 3.74-Å overall resolution.

#### (8) Δanchor *Tl*Sec

The initial set of 229,825 auto-picked particles were subjected to 2D classification, resulting in 105,578 particles. Three initial 3D models were generated by ab-initio reconstruction and refined by heterogenous refinement. One major class (76,726 particles) showed features of the Sec complex, and the particles from this class were used to generate the final map at 4.38-Å resolution with non-uniform refinement.

For additional Fourier shell correlation curves between the two half maps of final reconstructions, particle orientation distributions, local resolution distributions, see Supplementary Figs. 2-4.

### Atomic model building and generation of figures

Atomic models were built using Coot^53^. We first built models for *Sc*Sec[Sec62-] and ΔSec62 *Tl*Sec using our previous *Sc*Sec model (PDB ID 6N3Q; ref. ^28^) as a template. For ΔSec62 *Tl*Sec, we generated a homology model using SWISS-MODEL, which was rebuilt into the map using Coot. The *Sc*Sec[Sec62-] model was then used to build models for *Sc*Sec[C1] and *Sc*Sec[C2]. For *Sc*Sec62, a poly-alanine model was built into densities. Atomic models for all the mutant *Sc*Sec structures lacking Sec62 were also built starting from the *Sc*Sec[Sec62-] model. The *Sc*Sec[Sec62-] model was first fitted into each map using UCSF chimera and further fitted into the map in groups of domains and subunits using rigid-body refinement in Phenix (ref. ^54^). The models were then locally adjusted in Coot. Models for PM *Sc*Sec[C1] and PM *Sc*Sec[C2] were built similarly using the WT *Sc*Sec[C1] and *Sc*Sec[C2] models as starting models. Models for FN3mut ScSec[Sec62+] and PM/FN3mut *Sc*Sec[Sec62+] were built starting with the WT *Sc*Sec[C2] and PM/FN3mut *Sc*Sec[Sec62-] model, respectively. ΔSec62 *Tl*Sec was used as a starting model to build all *Tl*Sec structures.

The models were refined with Phenix real-space refinement using combined maps that were sharpened with a B-factor estimated based on the Guinier plot and low-pass-filtered at their overall resolution (produced by cryoSPARC). The refinement resolution was also limited to the overall resolution of the maps in Phenix. Secondary structure restraints were used during the refinement. MolProbity^55^ was used for structural validation. For refinement and validation statistics, see Tables 1 and 2.

UCSF Chimera^52^, ChimeraX^56^, and PyMOL (Schrödinger) were used to prepare figures in the paper. Unless stated otherwise, all shown cryo-EM maps are unsharpened maps that were low-pass-filtered at their overall resolution.

