## Supplementary Table 1 and Figures 1-4 for "Stepwise gating of the Sec61 protein-conducting channel by Sec63 and Sec62"

**Supplementary Table 1. Yeast strains and plasmids used in this study**

| Name | Genotype / Description | Source |
|---|---|---|
| **Yeast strains** | | |
| BY4741 | *MAT***a** *his3-1*, *leu2-0*, *met15-0*, *ura3-0* | Horizon Discovery |
| yMLT62 | *MAT***a** *leu2-0*::*pACT1-GEV*::*HIS3*, *rps9Δ*, *mek1Δ*, *his3-1*, *met15-0*, *ura3-0* | 47 |
| R1158 | BY4741 *URA3*::*pCMV-tTA* | 45 |
| TH_4087 | R1158 *pSEC61*::*KanMX*-tetO$_7$-*pCYC1* | 45 |
| TH_5187 | R1158 *pSEC63*::*KanMX*-tetO$_7$-*pCYC1* | 45 |
| ySI7 | BY4741 *Sbh1Δ*::*KanMX osw1Δ*::*HphMX*:: *SBH1*-15xGS-*SEC63*-TEV-GFP::*NatMX* | 28 |
| ySI8 | ySI7 *SEC61*(PM)*::LEU2* | This study |
| ySI42 | TH_5187 *SEC61*(PM)::*NatMX* | This study |
| ySI48 | TH_5187 *sbh1Δ*::*HphMX* | This study |
| ySI62 | R1158 *pSEC62*::*KanMX*-tetO$_7$-*pCYC1* | This study |
| ySI67 | yMLT62 *ura3-0*::*pGAL1-Tl*Sec::*NatMX* | This study |
| ySI73 | ySI48 *HO*::*SEC63*(E440R/F481S/440Δ7)::*LEU2* | This study |
| ySI74 | ySI42 *sbh1Δ*::*HphMX HO*::*SEC63*(E440R/F481S/440Δ7)::*LEU2* | This study |
| ySI77 | yMLT62 *ura3-0*:: *pGAL1-Tl*Sec(Δ*SEC62*)::*NatMX* | This Study |
| ySI89 | TH_4087 *SEC63*(E440R/F481S/440Δ7)::*NatMX*, | This study |
| ySI112 | ySI48 *HO*::*SEC63*(E440R/F481S/440Δ7/Δ210-216)::*LEU2* | This study |
| ySI113 | yMLT62 *ura3-0*::*pGAL1- Tl*Sec(Δanchor)::*NatMX* | This study |
| **Plasmids** | | |
| pYTK-001 | MoClo YTK part plasmid entry vector | 48 |
| pYTK-095 | MoClo YTK AmpR-ColE1 vector | 48 |
| pYTK-e101 | *URA3* integration vector containing a natMX expression cassette | This study |
| pYTK-e106 | *HO* integration vector containing a *LEU2* marker | This study |
| pYTK-e112 | CEN/ARS vector containing a *LEU2* marker | This study |
| pSI74 | pYTK-e106 *Sc*Sbh1-15xGS-*Sc*Sec63 (E440R/F481S/441Δ7)-TEV-GFP (with endogenous Sec63 promoter) | This study |
| pSI120 | pYTK-e106 *Sc*Sbh1-15xGS-*Sc*Sec63-FN3mut/Δ210-216-TEV-GFP (with endogenous Sec63 promoter) | This study |
| pSI123 | pYTK-e112 ScSec61 (with endogenous Sec61 promoter) | This study |
| pSI39 | pYTK-e112 *Sc*Sec62 (with endogenous Sec62 promoter) | This study |
| pSI5 | pYTK-e112 *Sc*Sec63 (with endogenous Sec63 promoter) | This study |
| pSI16 | pYTK-e112 *Sc*Sec63 (E440R/F481S/441Δ7) (with endogenous Sec63 promoter) | This study |
| pSI17 | pYTK-e112 *Sc*Sec63-TEV-GFP (with endogenous Sec63 promoter) | This study |
| pSI65 | pYTK-e101 *Tl*Sec (each subunit under a *GAL1* promoter) | This study |
| pSI87 | pYTK-e101 ΔSec62-*Tl*Sec (without Sec62) | This study |
| pSI94 | pYTK-e101 Δanchor-*Tl*Sec (with an anchor domain deletion mutant of Sec62) | This study |

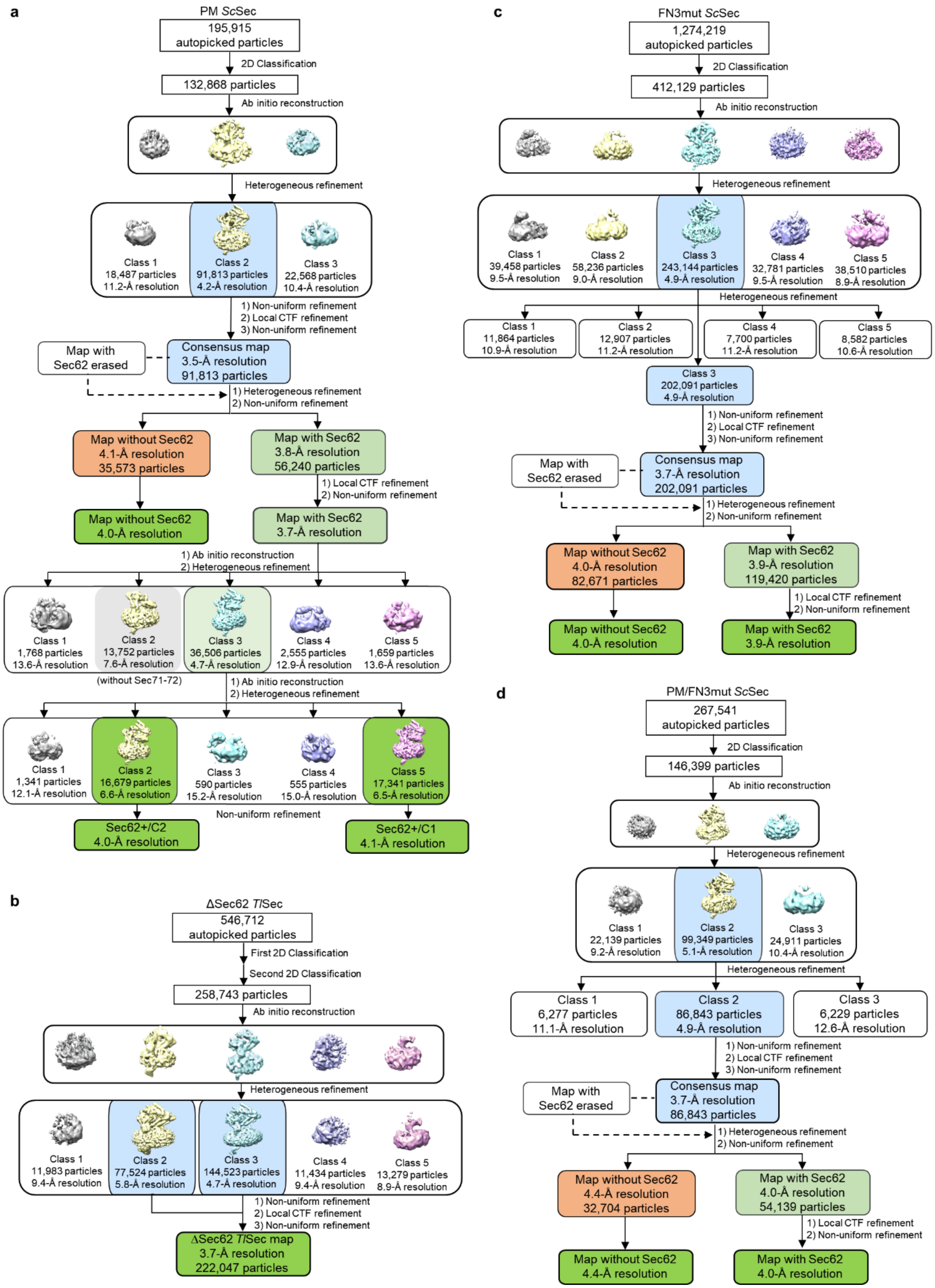


**Supplementary Figure 1. Cryo-EM image processing workflow**. Diagrams for single particle analysis procedure for pore mutant (PM) *Sc*Sec (**a**), ΔSec62 *Tl*Sec (**b**), E440R/F481S/441Δ7 (FN3mut) *Sc*Sec (**c**), and PM/FN3mut *Sc*Sec (**d**).

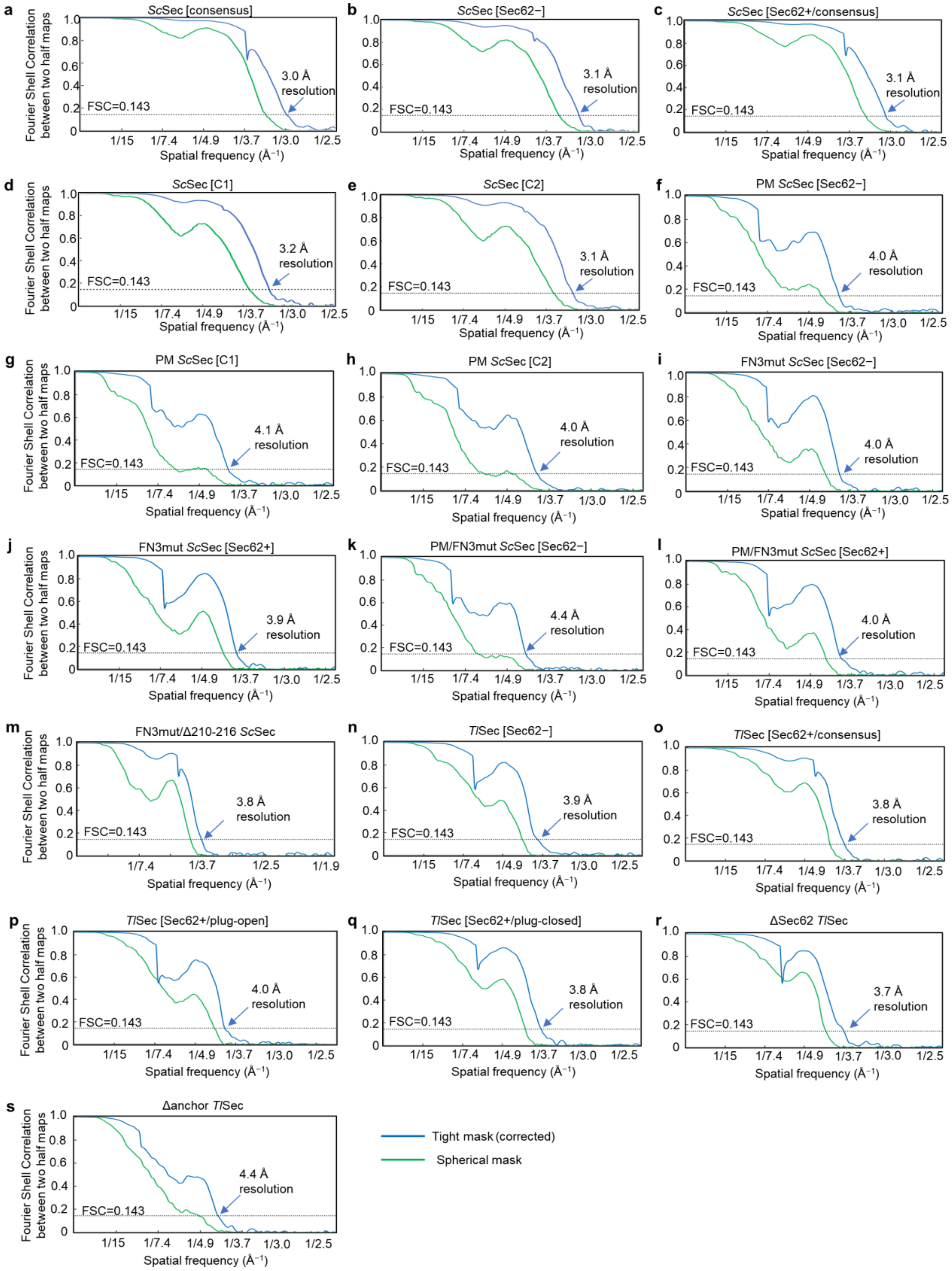


**Supplementary Figure 2. Fourier Shell Correlation (FSC) between two half maps of final 3D reconstructions**. FSC for maps derived from wildtype *Sc*Sec (**a**–**e**), pore mutant (PM) *Sc*Sec (**f-h**), E440R/F381S/441Δ7 (FN3mut) *Sc*Sec (**i**–**j**), PM/FN3mut *Sc*Sec (**k**–**l**), FN3mut/Δ210-216 *Sc*Sec (**m**), wildtype *Tl*Sec (**n-q**), ΔSec62 *Tl*Sec (**r**), and Δanchor *Tl*Sec (**s**).

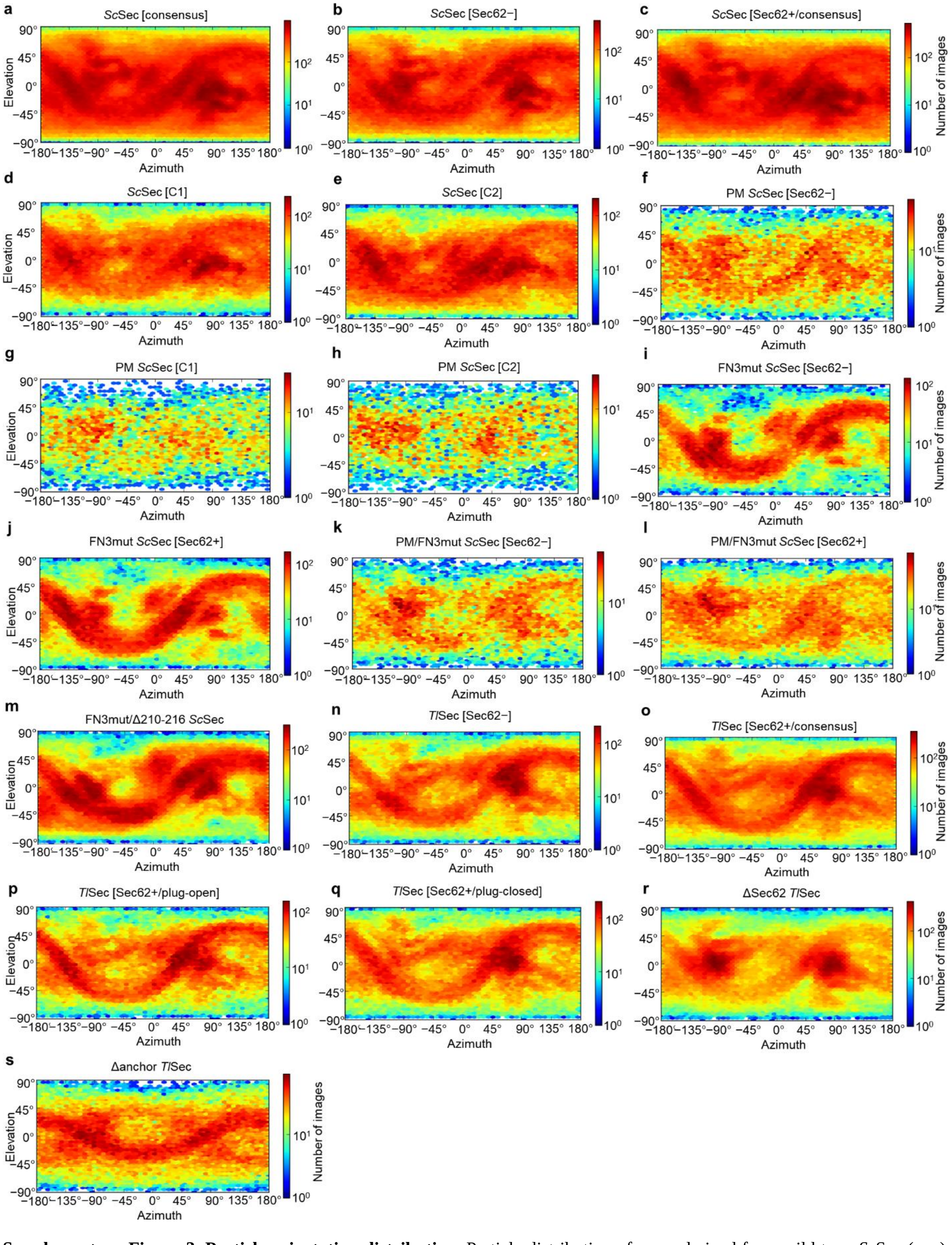


**Supplementary Figure 3. Particle orientation distribution**. Particle distribution of maps derived from wild type *Sc*Sec (**a**–**e**), pore mutant (PM) *Sc*Sec (**f-h**), E440R/F381S/441Δ7 (FN3mut) *Sc*Sec (**i**–**j**), PM/FN3mut *Sc*Sec (**k**–**l**), FN3mut/Δ210-216 *Sc*Sec (**m**), wild type *Tl*Sec (**n-q**), ΔSec62 *Tl*Sec (**r**), and Δanchor *Tl*Sec (**s**).

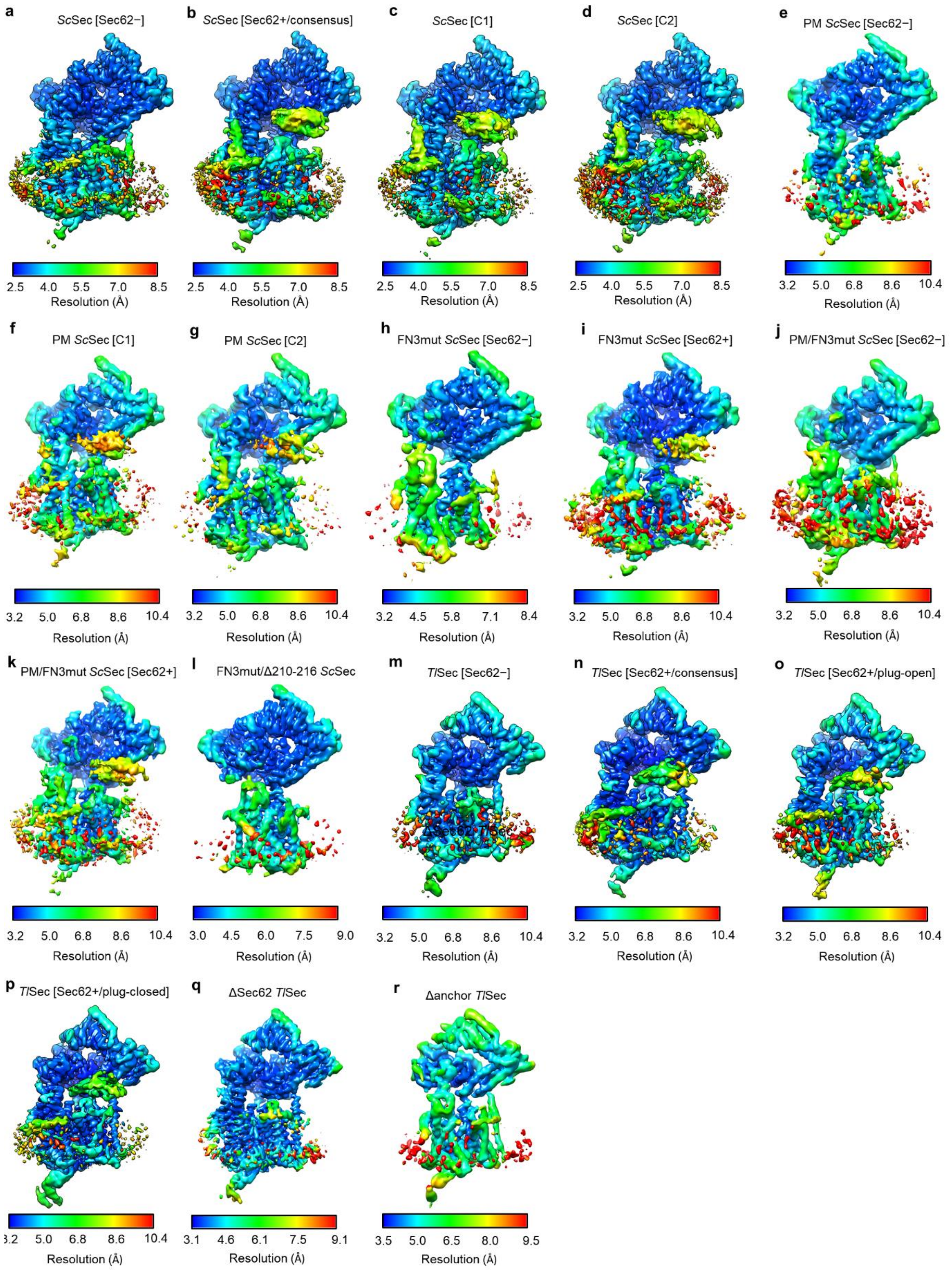


**Supplementary Figure 4. Local resolution distribution**. Local resolution distribution for maps derived from wild type *Sc*Sec (**a**–**d**), pore mutant (PM) *Sc*Sec (**e-g**), E440R/F381S/441Δ7 (FN3mut) *Sc*Sec (**h**–**i**), PM/FN3mut *Sc*Sec (**j**–**k**), FN3mut/Δ210-216 *Sc*Sec (**l**), wild type *Tl*Sec (**m-p**), ΔSec62 *Tl*Sec (**q**), and Δanchor *Tl*Sec (**r**).
